# Integrin-engaged Cellular Patches Mechanically Impose a Mitochondrial Respiratory Bottleneck to Suppress Cancer Cell Motility

**DOI:** 10.64898/2026.07.03.736349

**Authors:** Qizheng Zhang, Sona Roy, Tianrui Zhao, Wenjing Hou, Chenjie Xu, Jiancan Yu, Kai Wu, Xunwu Hu, Ye Zhang

## Abstract

The nanoscale organization of cell-adhesive ligands is increasingly recognized as a determinant of cell behavior, yet whether it directly regulates cellular metabolism remains unclear. Here we show that supramolecular clustering of integrin-binding ligands regulates mitochondrial respiratory capacity through integrin-mediated mechanotransduction. Supramolecular ligand clustering induces integrin redistribution and cytoskeletal remodeling, leading to mitochondrial reorganization and a selective constraint on oxidative phosphorylation. This respiratory limitation functionally constrains tumor cell migration and invasion and cannot be overcome by restoring cytoskeletal contractility, whereas replenishing mitochondrial metabolic substrates effectively rescues motility. In a HeLa xenograft model, the integrin-binding supramolecular system suppresses tumor growth and reduces extracellular matrix deposition. These findings identify mitochondrial respiratory capacity as a critical downstream effector of integrin mechanosignaling and establish extracellular ligand organization as a previously unrecognized driver of mechanically encoded metabolic regulation.

## Introduction

Metabolic reprogramming is a hallmark of cancer^1^ and has historically been framed by the Warburg effect^2^, which describes a preferential reliance on aerobic glycolysis^3^ rather than mitochondrial oxidative phosphorylation (OXPHOS)^4^ for ATP production. This conceptual framework motivated therapeutic strategies targeting glycolytic dependency^5, 6^. However, accumulating evidence has challenged this long-standing view^7^, revealing that cancer cells with high migratory and invasive capacities, including circulating tumor cell populations in murine and human models, can exhibit increased reliance on mitochondrial oxidative metabolism^8, 9^. Consistent with this view, clinical observations further underscore the relevance of mitochondrial respiration in tumor progression, as pharmacological inhibition of mitochondrial oxidative metabolism, such as with metformin, is associated with reduced relapse and improved outcomes in breast cancer patients^10, 11^. Collectively, these findings suggest that mitochondrial respiratory function is not merely a metabolic consequence of transformation, but a determinant of cellular fitness associated with aggressive tumor behavior. Importantly, mitochondrial respiration is closely linked to organelle morphological plasticity^12^. Dynamic balance between fission and fusion coordinates mitochondrial bioenergetic function^13^, whereas disruption of this equilibrium alters oxidative capacity and energetic homeostasis^14^. Given the substantial energetic demands imposed by migration and invasion^15^, mitochondrial structural dynamics may represent an important determinant of celullar adaptability in cancer^16, 17^.

Mitochondria are increasingly recognized as mechanosensitive organelles whose morphology is responsive to cytoskeletal forces^18–21^. Actin remodelling regulates mitochondrial fission-fusion dynamics and network architecture^22, 23^, establishing a physical and functional link between cytoskeletal organization and mitochondrial bioenergetic function. Integrins function as nanoscale sensors of extracellular matrix (ECM)^24–26^ organization, converting spatially defined ligand engagement into focal adhesion assembly, cytoskeletal organization and intracellular tension^27–29^. Previous studies have further established that nanoscale adhesive ligand presentation, including ligand clustering, spacing, and spatial order, regulates integrin organization, focal adhesion maturation, actin architecture, and cell migration. However, these studies have primarily defined canonical adhesion and cytoskeletal outputs^30^. Whether spatially organized integrin engagement can propagate beyond the cytoskeleton to regulate mitochondrial respiratory capacity and metabolic adaptability remains unclear. Building on this framework, we hypothesized that spatially programmed integrin engagement^31^ may regulate mitochondrial energetic function through mechanotransduction pathways and thereby influence migratory fitness. To test this hypothsis, we developed integrin-engaged cellular patches composed of self-assembling peptide nanofibrils that present integrin-binding ligands in defined supramolecular clusters (Figure 1a). This platform enables programmable control over integrin clustering and cytoskeletal organization^31, 32^, allowing us to examine how spatial encoding of integrin engagement influences mitochondrial bioenergetics and tumor cell behavior^33, 34^.

**Figure 1.**
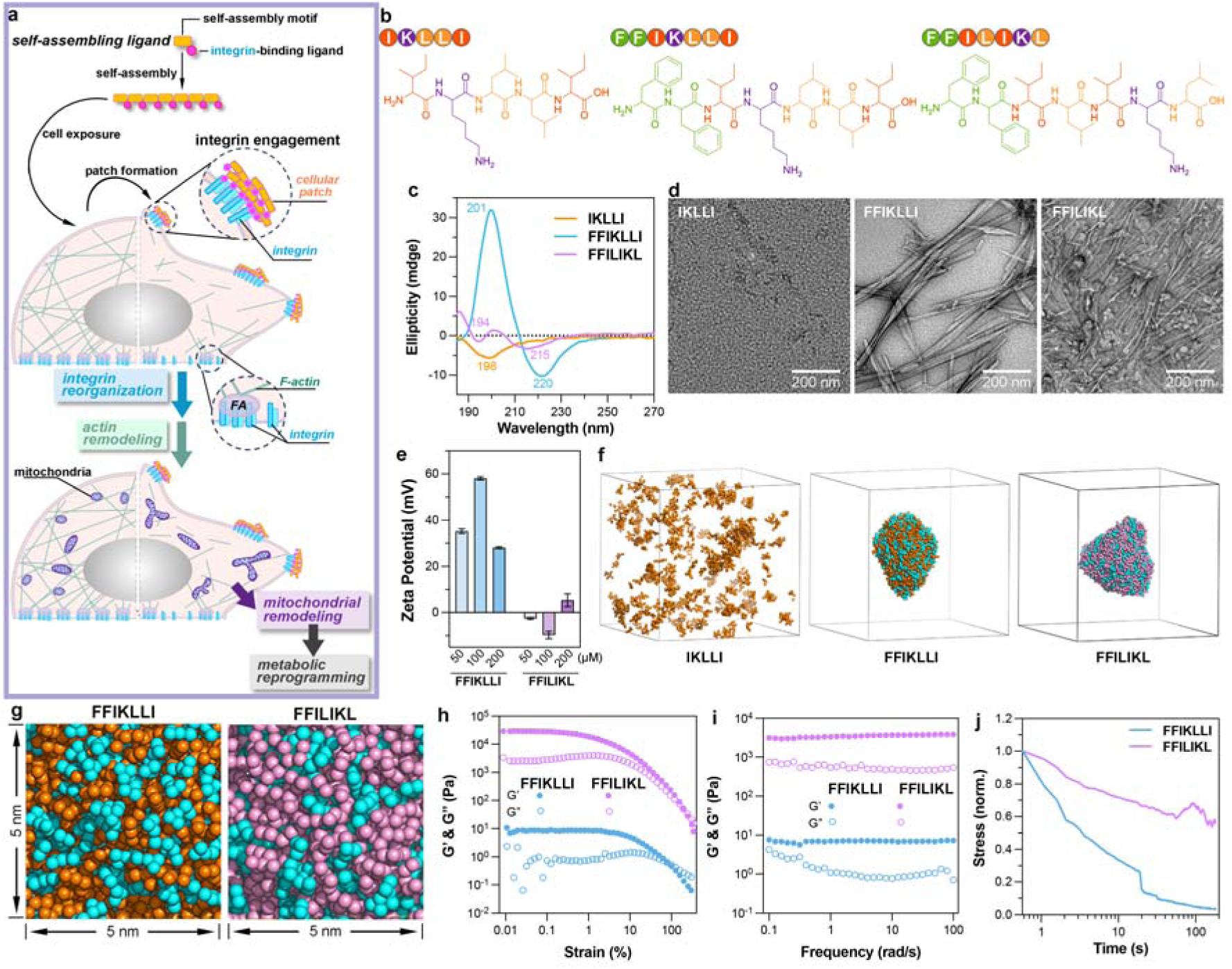
Engineering integrin ligands for self-assembly into molecularly ordered nanofibrils. **(a)** Schematic illustration of a self-assembling integrin ligand designed to regulate mitochondrial morphology via actin cytoskeleton remodeling, thereby suppressing metabolic constraints on cancer cell motility. **(b)** Chemical structures of laminin-derived integrin ligand **IKLLI**, self-assembling integrin ligand **FFIKLLI**, and the scrambled control sequece **FFILIKL**.**(c)** CD spectra of three peptides in water at a concentration of 10 mM. **(d)** TEM images of three peptides in water at a concentration of 10 mM. Scale bars represent 200 nm. **(e)** Zeta potentials of self-assemblies formed by **FFIKLLI** and **FFILIKL** in water at three concentrations (50, 100, and 200 μM). Data are presented as mean ± s.d. (*n* = 3). **(f)** Snapshots from coarse-grained molecular dynamics simulations of peptide self-assembly. Simulations were performed in a cubic box (21.5 nm per side) at a peptide concentration of 50 mM. Self-assembly motifs are shown in cyan, while integrin-binding motifs and their scrambled counterparts are shown in orange and magenta, respectively. Water molecules and counterions are omitted for clarity. **(g)** Representative snapshots of a 5 × 5 nm surface region extracted from the peptide self-assemblies shown in (f). **(h-j)** Rheological characterization of **FFIKLLI** and **FFILIKL** in water at a concentration of 10 mM. (h) Storage and loss moduli (G′ and G″) obtained from oscillatory strain sweep measurements at an angular frequency of ω = 6 rad/s. (i) G′ and G″ measured as a function of angular frequency during oscillatory frequency sweep experiments. (j) Stress relaxation behavior as a function of time under a constant applied strain at room temperature. Stress values were normalized to the initial stress recorded at the first time point.

## Results

### Engineering Integrin Ligands into Molecularly Ordered Nanofibrils that Form Cellular Patches

Integrin-mediated mechanotransduction is highly sensitive to the spatial organization of ligand presentation,^31, 35^ which governs integrin clustering and downstream force transmission^25, 36–38^. To create a programmable platform for controlling integrin organization, we coupled laminin-derived^39, 40^ *β*1 integrin-binding sequence **IKLLI**^41^ to an aromatic assembly motif capable of driving nanofibril formation through supramolecular interactions (Figure 1b). Our previous studies demonstrated that extending the assembly domain to three phenylalanines (FFF) generated highly stable fibrils that enforced persistent integrin clustering and cytoskeletal locking^31^. Here, we reduced the assembly domain to two phenylalanines (FF) to create a more dynamically reorganizable supramolecular platform while preserving molecular order. This design was motivated by emerging evidence that supramolecular dynamics can enhance biological responsiveness and cellular adaptation^42^. Structural and biophysical characterization confirmed that **FFIKLLI** assembled into molecularly ordered nanofibrils. CD spectroscopy, TEM imaging, and FTIR analysis demonstrated pronounced supramolecular ordering in **FFIKLLI** compared with free **IKLLI** and the scrambled control **FFILIKL** (Figure 1c,d and S1–S3). Zeta potential measurements (Figure 1e) together with coarse-grained molecular dynamics (MD) simulations (Figure 1f and S4) suggested preferential surface exposure and clustered presentation of **IKLLI** motifs, generating locally enriched adhesive patches that may facilitate multivalent integrin engagement (Figure 1g). Rheological analysis further indicated that **FFIKLLI** assemblies retained substantial molecular rearrangement, exhibiting lower stiffness and faster stress relaxation than **FFILIKL** networks (Figure 1h–j and S5). Together, these findings establish **FFIKLLI** as a molecularly ordered yet dynamically reorganizable supramolecular platform for spatially controlling integrin engagement.

To determine whether the supramolecular assemblies could form persistent cell-associated architectures, we examined their interactions with HeLa cells. Congo Red staining and scanning electron microscopies (SEM) revealed that both **FFIKLLI** and the scrambled control **FFILIKL** formed membrane-associated supramolecular assemblies (Figure 2a-e, S6, and S7). Notably, **FFIKLLI** generated extended nanofibrillar networks that preferentially accumulated at cell edges and closely associated with membrane protrusions, forming spatially organized cell-associated patches (Figure 2d). In contrast, **FFILIKL** formed compact aggregates with limited fibrillar organization and a diffuse surface distribution (Figure 2e). Both **IKLLI** and **FFIKLLI** maintained high cellular viability across the tested concentrations, whereas **FFILIKL** reduced viability at higher doses (Figure 2f). These findings demonstrate that sequence-directed self-assembly enables **FFIKLLI** to form cytocompatible nanofibrillar patches at the cell surface, positioning integrin-binding ligands within organized adhesive interfaces for subsequent cell signaling.

**Figure 2.**
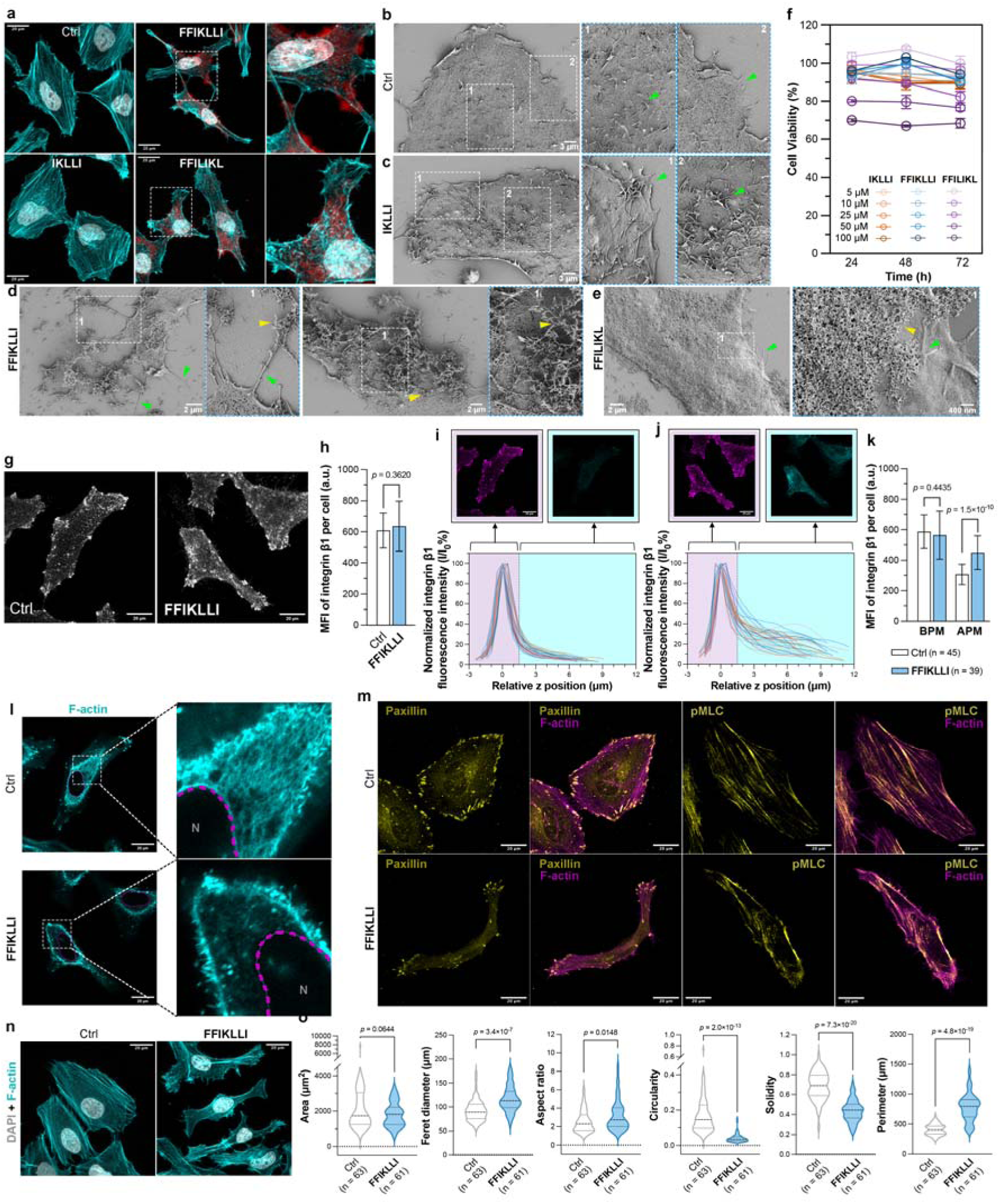
Cell-associated nanofibrillar patches redistribute activated β1 integrin and remodel cytoskeletal organization. **(a)** Maximum-intensity projection (MIP) images of HeLa cells with or without peptide treatment (100 μM, 24 h), co-stained with phalloidin (cyan), DAPI (gray), and Congo Red (red). **(b-e)** SEM images of HeLa cells with or without peptide treatment (100 μM, 24 h). Scale bars represent 3 μm in (b, c) and 2 μm in (d, e). Enlarged views highlight the cell edge and apical plasma membrane, with green arrows denoting intact filopodia/microvilli, and yellow arrows denoting those entangled or covered by self-assembling nanofibrils. **(f)** HeLa cell viability following peptide treatment at the indicated concentrations for 3 days. **(g)** MIP images of activated integrin β1 immunofluorescence in control and **FFIKLLI**-treated cells (100 μM, 24 h). **(h)** Mean fluorescence intensity (MFI) of active integrin β1 per cell, corresponding to (g). Data present mean ± s.d. (*n* = 45 and 39 cells for control and **FFIKLLI**-treated, respectively). Two-tailed Student’s *t*-test. **(i,j)** Z-axis distribution of active integrin β1 from basal to apical membranes in control (*n* = 45) and **FFIKLLI**-treated (*n* = 39) cells. The paxillin-defined focal adhesion plane was set as z = 0 and used as the reference intensity (I_₀_), with fluorescence at each position expressed as I/I_₀_. Integrin stacks spanning the basal (−3–1.5 μm, magenta) and apical (1.5–12 μm, cyan) regions are shown in the corresponding colors. **(k)** Quantification of active integrin β1 fluorescence per cell at the basal and apical plasma membranes. Data represent mean ± s.d. Two-tailed Student’s t-test. **(l)** Representative F-actin (cyan) images showing a central cytoplasmic z-slice of the stack, selected to exclude the basal focal adhesion layer. Nuclei (N) are outlined with magenta dashed lines. **(m)** Immunofluprescence of focal adhesion (paxillin, yellow) and contractility (pMLC, cyan) with phalloidin (magenta) in HeLa cells treated with or without 100 μM **FFIKLLI** for 24 h. **(n)** Representative images of HeLa control and **FFIKLLI**-treated (100 μM, 24 h) cells co-stained with phalloidin (cyan) and DAPI (gray). **(o)** Quantification of cell morphology under the conditions shown in (n). Violin plots display the median and quartiles. Two-tailed Welch’s t-test. Scales bars in (a, g, i, j, l, m and n) represent 20 μM.

### Molecularly Ordered Nanofibrillar Patches Reorganize Integrin Signaling and Suppress Tumor Cell Motility

Having established that **FFIKLLI** nanofibrils form membrane-associated assemblies on cells, we next examined how these interfaces influence integrin organization and downstream cytoskeletal remodeling. Immunofluorescence imaging of activated integrin *β*1 revealed no significant difference in overall integrin activation between control and **FFIKLLI**-treated cells (Figure 2g and 2h). However, z-resolved analysis uncovered a pronounced redistribution of activated integrin *β*1 from the basal focal adhesion plane toward the apical membrane following **FFIKLLI** treatment (Figure 2i–k, S8), resulting in reduced basal and enhanced apical signals. These findings suggest that nanofibrillar patches spatially redistribute activated integrin *β*1 without increasing overall integrin activation. Notably, the same cells exhibited substantial alterations in actin organization. F-actin staining revealed disrupted filament architecture and loss of stress-fiber integrity in **FFIKLLI**-treated cells compared with controls (Figure 2l and S9), indicating that spatial redistribution of activated integrin *β*1 is accompanied by extensive cytoskeletal remodeling. To determine whether this altered integrin organization and cytoskeletal remodeling propagate into downstream mechanotransduction pathways, focal adhesion assembly and actomyosin contractility were evaluated (Figure 2m and S10). **FFIKLLI** treatment markedly reduced paxillin-positive focal adhesions compared with untreated cells, indicating impaired focal adhesion maturation. Consistent with diminished adhesion-dependent signaling, phosphorylation of myosin light chain (pMLC) was substantially decreased following **FFIKLLI** treatment, suggesting attenuation of actomyosin-generated tension.

At the cellular level, these changes in cytoskeletal organization and contractility were reflected in a distinct morphological phenotype. Quantitative morphometric analysis showed increased Feret diameter and aspect ratio together with reduced circularity and solidity (Figure 2n and 2o), indicating a more elongated and less compact cellular morphology. These observations further support widespread remodeling of cytoskeletal architecture and cellular mechanical state following **FFIKLLI** treatment. Marked reductions in cell migration and invasion accompanied these structural changes. Wound-healing assays demonstrated that **FFIKLLI** suppressed HeLa cell migration in a concentration-dependent manner, whereas **IKLLI** produced no detectable effect and the scrambled control **FFILIKL** induced only modest inhibition (Figure 3a and S11). Consistently, Transwell invasion assays revealed a similar trend, with **FFIKLLI** producing the strongest suppression of invasive behavior (Figure 3b and 3c). Collectively, these findings demonstrate that molecularly ordered **FFIKLLI** nanofibrils spatially reorganize activated integrin *β*1, remodel actin architecture, impair focal adhesion maturation and actomyosin contractility, and are associated with marked suppression of tumor cell migration and invasion.

**Figure 3.**
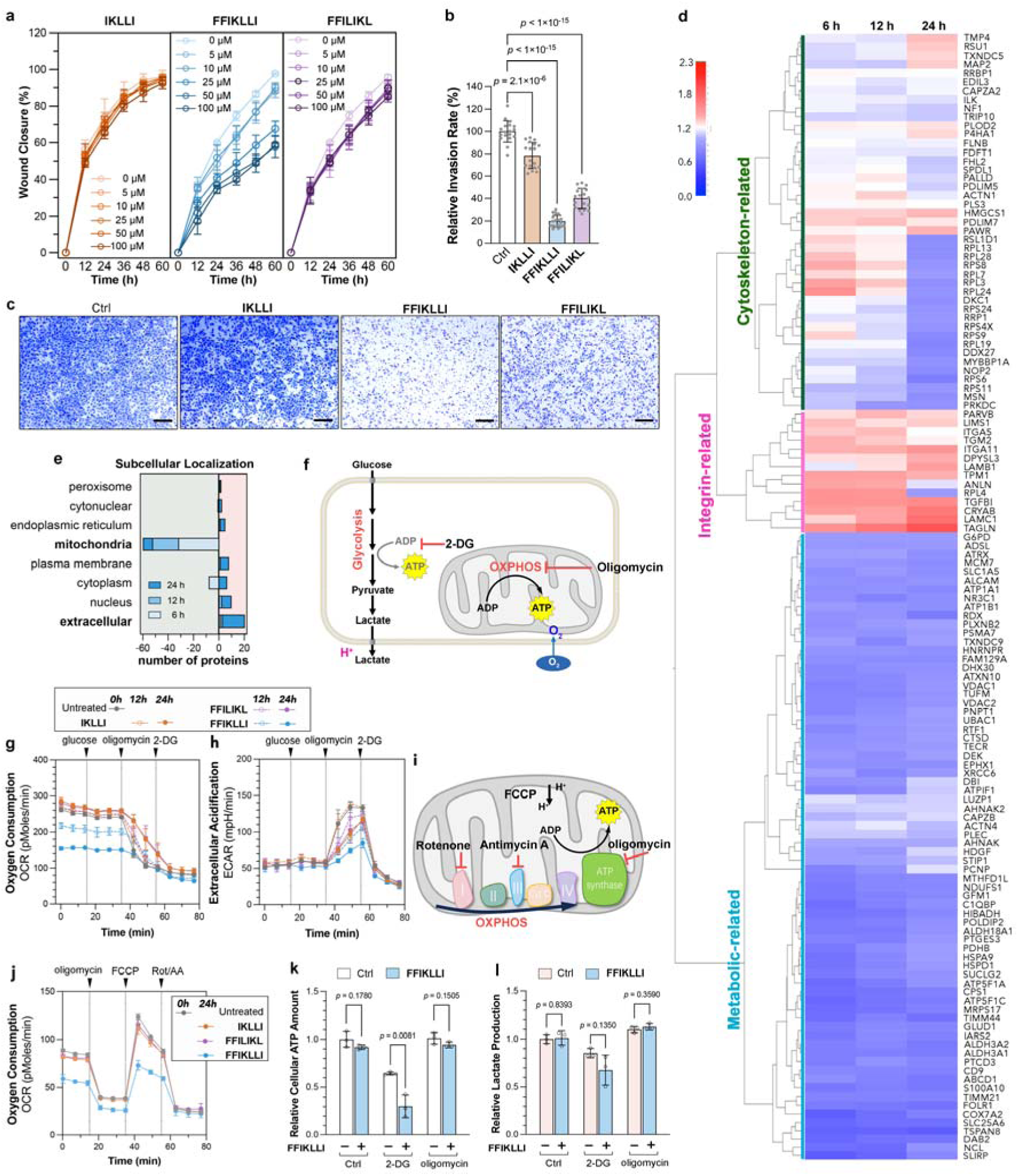
Integrin-binding FFIKLLI assemblies reprograms cellular metabolism and constrain oxidative phosphorylation. **(a)** HeLa cell wound closure rates following peptide treatment at the indicated concentrations. **(b,c)** Quantification of invasive HeLa cells with or without the indicated treatments (b) and the representative bright-field images of crystal violet-stained invasive HeLa cells under the corresponding conditions (c). Statistical significance was assessed using Brown-Forsythe and Welch ANOVA followed by Dunnett’s T3 post-hoc comparisons versus the control group. Scale bars, 200 μm. **(d)** Heatmap showing the relative abundance of cytoskeleton-, integrin- and metabolismp-associated proteins in HeLa cells treated with **FFIKLLI** (50 μM) for 6, 12 and 24 h relative to untreated controls. **(e)** Gene Ontology (GO) cellular component enrichment of differentially expressed proteins across time points. **(f)** Schematic of glycolysis and oxidative phosphorylation (OXPHOS) pathways, indicating sites of inhibition by 2-deoxy-D-glycose (2-DG) and olygomcin for metabolic flux measurements. **(g, h)** Oxygen consumption rate (OCR, g) and extracellular acidification rate (ECAR, h) of HeLa cells treated with peptides (100 μM) for 12 and 24 h, with sequential injection of glucose, oligomycin and 2-DG to asssess glycolytic and mitochondrial activity. **(i)** Schematic illustration of the principle underlying the mitochondrial stress test. **(j)** OCR traces following peptide treatment (100 μM, 12 h) with sequential injection of oligomycin, FCCP, and rotenone/antimycin A (Rot/AA) to evaluate mitochondrial respiration.. Group differences were analyzed using Brown–Forsythe and Welch ANOVA followed by Dunnett’s T3 post-hoc test versus untreated controls; adjusted p values are indicated. **(k, l)** ATP levels (k) and lactate production (l) in HeLa cells treated with **FFIKLLI** (100 μM, 24 h) under glycolysis inhibition (2-DG) or OXPHOS inhibition (oligomycin). Values are normalized to untreated, non-inhibited controls. Two-tailed Student’s t-test was used for statistical comparisons.

### Integrin-Engaged Nanofibrillar Patches Suppress Mitochondrial Respiratory Capacity

The marked suppression of cell migration and invasion prompted us to investigate whether **FFIKLLI**-induced mechanotransduction influences mitochondrial energy metabolism. To obtain a systems-level view of cellular responses, HeLa cells were treated with **FFIKLLI** and subjected to quantitative proteomic analysis. Quantitative proteomic profiling revealed coordinated remodeling of cellular metabolism following **FFIKLLI** treatment (Figure 3d). Gene Ontology analysis showed preferential downregulation of mitochondrial-associated proteins together with enrichment of extracellular matrix- and adhesion-related components (Figure 3e and Figure S12, S13). Consistent with these changes, KEGG pathway enrichment highlighted oxidative phosphorylation and the tricarboxylic acid (TCA) cycle among the pathways significantly enriched in the downregulated proteome (Figure S14), while protein domain enrichment analysis identified the mitochondrial carrier domain as the most significantly enriched downregulated domain (Figures S15). Together, these orthogonal analyses consistently support coordinated remodeling of extracellular adhesion programs and mitochondrial metabolic machinery following **FFIKLLI** treatment.

To directly evaluate cellular energy metabolism, we monitored oxygen consumption rate (OCR) and extracellular acidification rate (ECAR) during a glycolysis stress-test sequence involving glucose, oligomycin, and 2-deoxy-D-glucose (2-DG) (Figure 3f). **FFIKLLI** treatment induced a pronounced and time-dependent reduction in OCR (Figure 3g), whereas ECAR was only modestly affected (Figure 3h). These findings suggested a preferential impairment of mitochondrial respiration and prompted a more detailed assessment of respiratory capacity. To further define mitochondrial energetic capacity, we performed a Seahorse mitochondrial stress test by sequentially injecting oligomycin, FCCP, and rotenone/antimycin A (Rot/AA) (Figure 3i). **FFIKLLI**-treated cells exhibited reduced basal respiration relative to untreated and control peptide-treated cells, indicating suppression of steady-state mitochondrial activity (Figure 3j and S16). Following ATP synthase inhibition, OCR remained consistently lower in **FFIKLLI**-treated cells, demonstrating diminished ATP-linked respiration. FCCP-induced maximal respiration was also markedly blunted, resulting in a substantial reduction in spare respiratory capacity. In contrast, Rot/AA reduced OCR to comparable minimal levels across all groups, confirming that non-mitochondrial oxygen consumption remained unaffected. Collectively, these findings demonstrate that integrin-directed supramolecular assemblies impose a persistent limitation on mitochondrial respiratory function at both basal and maximal respiratory states.

Despite the marked reduction in mitochondrial respiration, ATP levels were only modestly decreased under basal conditions (Figure 3k), suggesting compensatory maintenance of cellular energy homeostasis. However, inhibition of glycolysis with 2-DG resulted in substantially greater ATP depletion in **FFIKLLI**-treated cells than in control cells, indicating compensatory reliance on glycolytic ATP production when mitochondrial respiratory capacity is restricted. In contrast, oligomycin produced only a minor additional decrease in ATP levels, consistent with a pre-existing limitation in mitochondrial ATP production. Correspondingly, lactate production remained largely unchanged across conditions (Figure 3l), suggesting preservation of glycolytic activity despite impaired mitochondrial respiration. Together, these findings indicate that integrin-engaged nanofibrillar patches do not induce a generalized metabolic collapse but instead impose a selective limitation on mitochondrial respiratory capacity while preserving glycolytic compensation. Consequently, cellular ATP homeostasis and cell viability remain largely intact uncer basal conditions. However, the loss of respiratory flexibility may become functionaly limiting when cells encounter energetically demanding processes such as migration and invasion.

### Mitochondrial Respiratory Limitation Underlies the Anti-migratory Effect of Integrin-Engaged Cellular Patches

Given the close relationship between mitochondrial architecture and respiratory function, we examined whether **FFIKLLI**-induced metabolic remodeling was accompanied by alterations in mitochondrial organization. Time-lapse live-cell imaging demonstrated progressive and coordinated remodeling of both the actin cytoskeleton and mitochondrial architecture following **FFIKLLI** treatment. As cells gradually adopted an elongated morphology, mitochondria underwent concomitant redistribution and elongation, forming extended networks aligned with the remodeled cytoskeletal architecture (Figure 4a and S17). Quantitative morphometric analysis confirmed a significant shift from punctate to filamentous mitochondrial populations (Figure 4b-d, and S18-20), indicating extensive mitochondrial network remodeling. Notably, mitochondrial membrane potential remained largely unchanged throughout treatment, whereas the uncoupler CCCP markedly reduced membrane polarization (Figure 4e and 4f), suggesting that mitochondrial integrity was preserved despite substantial architecture reorganization.

**Figure 4.**
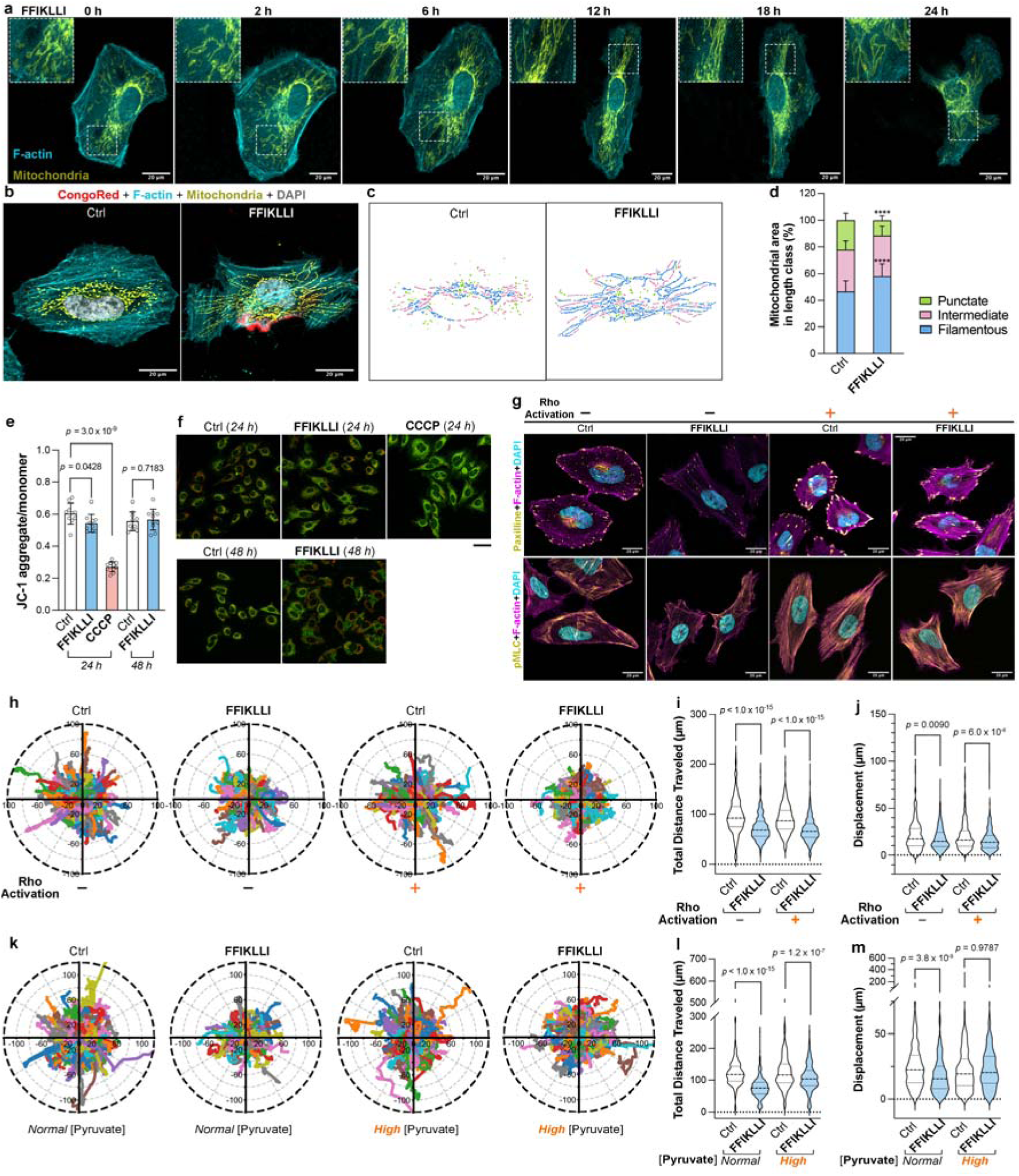
Mitochondrial network remodeling links integrin-directed cytoskeletal reprogramming to metabolic control of cell migration. **(a)** Time-lapse live-cell imaging of HeLa cells co-expressing mEGFP-LifeAct (F-actin, cyan) and DsRed2-Mito (mitochondria, yellow) following treatment with 100 μM **FFIKLLI**. (**b**) TOM20 immunofluorescence (yellow) with phalloidin (cyan), DAPI (gray) and Congo Red (red) in HeLa cells treated with or without 100 μM **FFIKLLI** for 24 h. Scale bar, 20 μm. (**c**,**d**) Quantification of mitochondrial morphology corresponding to (a), classifying mitochondria per cell as punctate (green), intermediate (pink), or filamentous (blue).Data are compositional and presented as mean ± s.d.; *n* = 63 and 61 cells. Statistical significance was assessed using the Hotelling-Lawley trace from multivariate analysis of variance (MANOVA) on centered log-ratio (CLR)-transformed data. **** represents *p* ≤ 0.0001. (**e,f**) Quantification and representative images of mitochondrial membrane potential assessed by JC-1 in HeLa cells treated with or without 100 μM **FFIKLLI** for 24 or 48 h. CCCP-treated cells were used as a positive control. Data are mean ± s.d; *n* = 10 fields of view. Statistical significance was assessed using two-tailed Welch’s *t*-test. Scale bar, 50 μm. (**g**) Paxillin and pMLC immunofluorescence (yellow) in HeLa cells treated with or without 100 μM **FFIKLLI** for 24 h followed by Rho activator II treatment (4 h) where indicated. Cells were co-stained with phalloidin (magenta) and DAPI (cyan). Scale bar, 20 μm. (**h-m**) Trajectory plots and migration analysis showing total travel distance and displacement for cells pretreated with or without **FFIKLLI** (100 μM, 24 h). Cells were subsequently treated with Rho activator II treatment (2 h) where indicated (h-j), or analysed under normal (1 mM) or high (5 mM) pyruvate conditions (k-m). *n* = 314, 374, 254, 467 and 447, 382, 375, 480 cells, respectively. Statistical significance was assessed using two-tailed Student’s *t*-test.

Pharmacological activation of Rho signaling restored both focal adhesion formation and myosin phosphorylation (Figure 4g, and S21-23), demonstrating substantial restoration of focal adhesion signaling and actomyosin contractility. Surprisingly, recovery of actomyosin contractility failed to rescue cell motility. Single-cell tracking revealed that **FFIKLLI**-treated cells exhibited substantially reduced migration distance and net displacement, and these defects persisted despite Rho activation (Figure 4h–j). These findings indicate that restoration of cytoskeletal contractility is insufficient to recover cell motility, suggesting that additional downstream mechanisms contribute to the migratory defect. In contrast to Rho-mediated recovery of contractility, increasing extracellular pyruvate largely rescued both migration distance and net displacement in **FFIKLLI**-treated cells (Figure 4k–m), indicating that mitochondrial respiratory capacity, rather than mechanical tension alone, is a dominant determinant of migratory competence under these conditions. Consistent with this functional recovery, pyruvate treatment also largely restored mitochondrial network morphology toward the control state (Figure S24, S25), supporting a close association between mitochondrial structural remodeling and migratory competence. Together, these findings demonstrate that suppression of cell motility cannot be explained solely by altered cytoskeletal mechanics. Instead, **FFIKLLI**-induced integrin reorganization establishes a mitochondrial respiratory. The concordant restoration of mitochondrial network organization and cell motility following pyruvate supplementation further supports mitochondrial function as a critical downstream effector of integrin-mediated mechanotransduction.

### Integrin-directed Supramolecular Assemblies Constrain Tumor Growth and Matrix Remodeling In Vivo

To determine whether the mitochondrial respiratory limitation imposed by **FFIKLLI** affects long-term cellular fitness, we evaluated cell viability during prolonged exposure (Figure 5a). Concentrations up to 200 μM exerted minimal effects on viability over several days, whereas higher concentrations (500 μM) induced a gradual decline in survival, reaching approximately 60% viability after 7 days. These findings indicate that **FFIKLLI** does not induce acute cytotoxicity but instead establishes a persistent metabolic state that remains compatible with short-term survival while progressively llimiting long-term cellular fitness.

**Figure 5.**
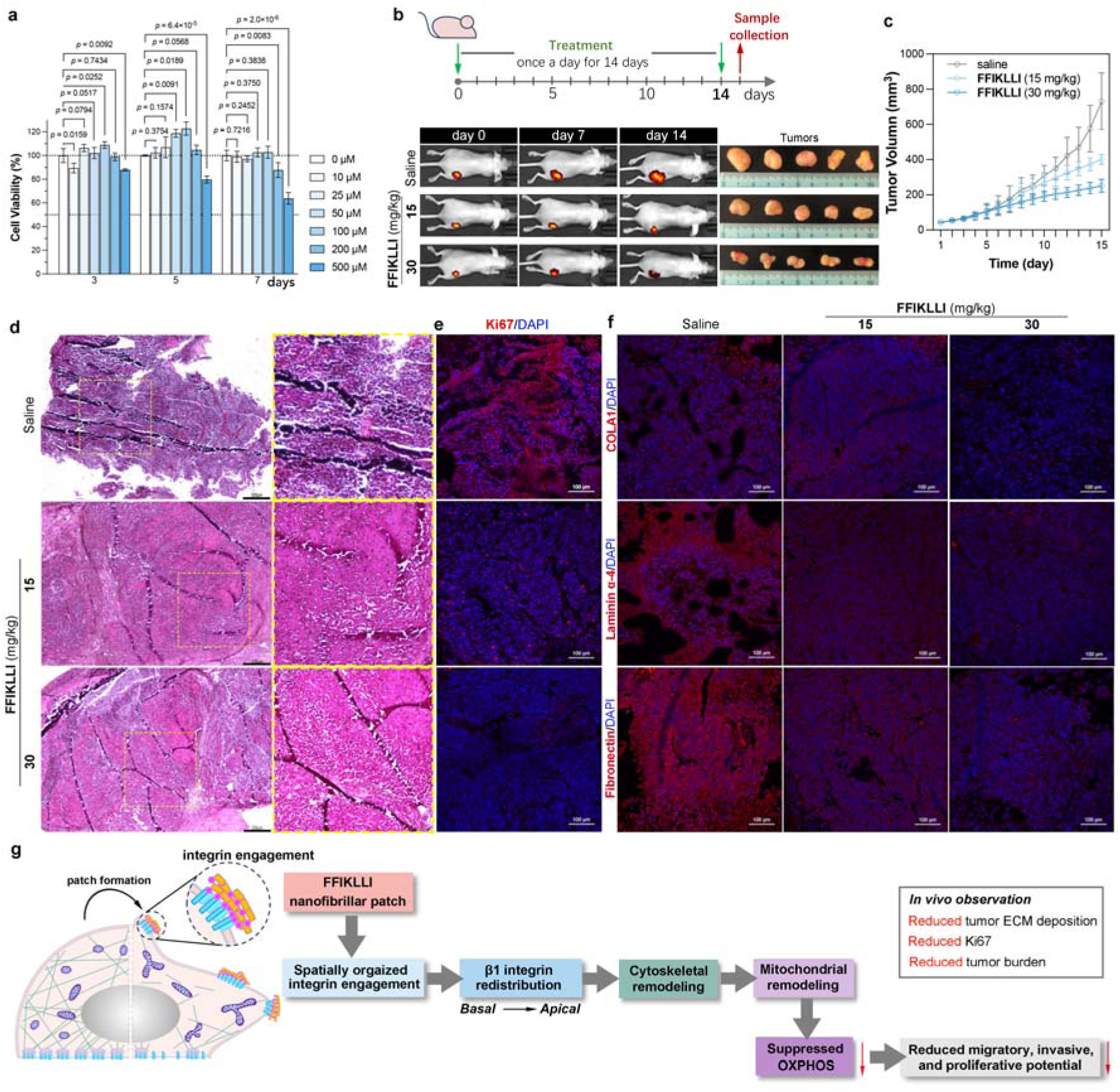
Peritumoral administration of the FFIKLLI suppresses tumor growth and remodels ECM in a HeLa xenograft model. **(a)** Long-term (7 days) cell viability of HeLa cells treated with **FFIKLLI** at indicated concentrations. (**b**,**c**) Tumor growth in HeLa xenografts following daily peritumoral injections of saline or FFIKLLI (15 or 30 mg/kg). In vivo fluorescence imaging (b) was performed every 7 days, and tumor volumes were measured by calipers (c) throughout treatment. Representative tumor photographs of excised tumor at day 15 (b) are shown. Data are mean ± s.d. (n = 5). (**d**) Representative hematoxylin and eosin (H&E) staining of tumor sections. Scale bar, 200 μm. (**e**,**f**) Immunofluorescence staining of tumor sections. (e) Ki67 (red) with DAPI (blue). (f) ECM adhesion proteins—collagen I α1 (COL1A1), laminin α4, and fibronectin (red)—with DAPI (blue). Scale bars, 100 μm. (**g**) Schematic illustration of **FFIKLLI** self-assembly engaging integrin clusters at the apical plasma membrane, triggering cytoskeletal reorganization, limiting mitochondrial OXPHOS, diminishing adhesion strength, and inhibiting tumor cell migration and invasion.

To evaluate the *in vivo* consequences of integrin-directed supramolecular regulation, we examined **FFIKLLI** in a HeLa xenograft model. Peritumoral administration of **FFIKLLI** once daily for 14 days produced dose-dependent suppression of tumor growth compared with vehicle-treated controls (Figure 5b), without significant changes in body weight (Figure S26), indicating minimal systemic toxicity. Consistent with these observations, both tumor volume and excised tumor weight were progressively reduced with increasing dose and treatment duration (Figures 5c, S27). Histological examination by hematoxylin and eosin (H&E) staining revealed reduced tumor cellularity (Figure 5d), while Ki67/DAPI co-staining demonstrated a dose-dependent decrease in proliferative index, indicating suppression of tumor growth *in vivo* (Figures 5e, S28a).

Because integrin-mediated signaling regulates not only intracellular mechanics but also reciprocal interactions with the extracellular matrix, we next examined whether the tissue microenvironment was altered following treatment. Immunofluorescence staining revealed a dose-dependent reduction in collagen I, laminin α4, and fibronectin following **FFIKLLI** treatment (Figure 5f, S28b-d). These findings indicate that integrin-directed supramolecular regulation influences not only tumor growth but also the composition of the surrounding extracellular matrix. The coordinated reduction in collagen I, laminin α4, and fibronectin is consistent with attenuation of a tumor-supportive extracellular microenvironment, suggesting that the cellular reprogramming induced by **FFIKLLI** extends beyond intracellular signaling to influence tissue-level matrix organization. Together, these findings demonstrate that integrin-engaged supramolecular assemblies suppress tumor progression across multiple biological scales, from intracellular mechanotransduction and mitochondrial metabolism to tissue-level extracellular matrix organization (Figure 5g). By spatially organizing integrin engagement through molecularly ordered cellular patches, FFIKLLI establishes a persistent limitation in mitochondrial respiratory capacity that restricts tumor cell molitily and ultimately constrains tumor growth *in vivo*.

## Discussion

We identify mitochondrial respiratory capacity as a functional downstream output of integrin-mediated mechanotransduction and further demonstrate that spatial organization of extracellular adhesive cues is sufficient to initiate this mechanometabolic response. By spatially clustering integrin-binding ligands within supramolecular cellular patches, we show that adhesive architecture can influence cellular bioenergetics independently of changes in overall integrin activation. Rather than globally enhancing integrin signaling, supramolecular ligand organization redistributed activated integrin *β*1 and reorganized downstream cytoskeletal architecture, ultimately constraining mitochondrial respiratory capacity. Notably, restoration of cytoskeletal contractility failed to recover migratory behavior, whereas replenishment of mitochondrial metabolic substrates largely restored motility. Together, these findings identify mitochondrial respiratory capacity as a key functional effector through which extracellular adhesive organization regulates cell behavior.

Whereas previous studies have established that intracellular mechanical states influence cellular metabolism^43^, our findings further demonstrate that extracellular adhesive organization itself is sufficient to initiate this mechanometabolic program. Accordingly, Our findings position extracellular adhesive architecture as an initiating extracellular determinant of a mechanotransductive cascade linking extracellular organization to mitochondrial respiration function. Although mitochondria are increasingly recognized as mechanosensitive organelles, the mechanisms linking extracellular organization to mitochondrial function remain incompletely understood.

The observation that mitochondrial remodeling occurred without substantial changes in mitochondria-associated DRP1 levels indicates that pathways beyond canonical fission regulation may contribute to this response (Figure S29, S30). These observations suggest that mitochondrial remodeling and respiratory suppression are coordinated downstream consequences of integrin-mediated mechanotransduction, although the causal relationship between mitochondrial architecture and respiratory capacity remains to be determined. Beyond its effects on mitochondrial respiration and cell motility, integrin-directed supramolecular organization was also associated with reduced deposition of collagen I, laminin α4, and fibronectin in tumor tissues, consistent with the established role of integrin signaling in coordinating cell–matrix interactions. Together, these findings establish extracellular ligand organization as a previously underappreciated upstream regulator of mitochondrial respiratory capacity and suggest that spatial organization within the extracellular microenvironment constitutes an instructive mechanical cue through which cellular energetic state and behavior are coordinated.

## Materials and methods

### Peptide synthesis

Peptides **IKLLI**, **FFIKLLI**, **FFILIKL** were custom-synthesized by solid-phase synthesis with a purity of ≥ 95% (GL Biochem, China). The structures of **IKLLI** and **FFIKLLI** were previously reported and validated by NMR and LC-MS in our earlier work. Characterization data for the remaining peptides are provided below.

### FFILIKL

^1^H NMR (500 MHz, DMSO-*d*_6_) *δ* 8.63 (s, 1H), 8.29 (d, *J* = 8.3 Hz, 1H), 8.14 (d, *J* = 8.0 Hz, 1H), 8.06 (d, *J* = 7.7 Hz, 1H), 7.96 (d, *J* = 7.8 Hz, 1H), 7.82 – 7.58 (m, 3H), 7.34 – 7.14 (m, 10H), 4.71 (s, 1H), 4.38 (dd, *J* = 14.9, 8.0 Hz, 1H), 4.28 – 4.21 (m, 3H), 4.21 – 4.16 (m, 1H), 3.90 (s, 1H), 3.07 (d, *J* = 13.8 Hz, 1H), 2.98 (d, *J* = 10.7 Hz, 1H), 2.87 (dd, *J* = 12.2, 8.4 Hz, 1H), 2.79 (dd, *J* = 13.8, 9.6 Hz, 1H), 2.76 – 2.71 (m, 2H), 1.71 (s, 2H), 1.67 – 1.56 (m, 3H), 1.56 – 1.37 (m, 9H), 1.30 (s, 2H), 1.13 – 1.03 (m, 2H), 0.88 (d, *J* = 6.5 Hz, 6H), 0.86 – 0.77 (m, 18H). ^13^C NMR (126 MHz, DMSO-*d*_6_) *δ* 173.81, 171.42, 171.09, 170.57, 170.40, 170.34, 137.30, 129.52, 129.13, 128.31, 127.97, 126.91, 126.28, 56.71, 56.41, 53.60, 51.81, 50.85, 49.95, 40.55, 38.62, 37.55, 36.71, 36.50, 31.25, 26.52, 24.24, 24.09, 24.04, 23.95, 23.00, 22.80, 21.91, 21.50, 21.06, 15.15, 15.03, 10.87. LC-MS (ESI) for C_48_H_76_N_8_O_8_, [M+H]^+^ calcd 893.59, found 893.65 (*m*/*z*).

### Preparation of peptide assemblies

**IKLLI** and **FFIKLLI** were dissolved in sterile-filtered Milli-Q water (0.22 μm PES filter, Sartorius) at 10 mM and sonicated for 30 min at room temperature. The pH was adjusted to 7.0 using 1 M NaOH. Owing to its limited aqueous solubility, **FFILIKL** was initially dissolved in DMSO and subsequently diluted with water to obtain a 10 mM stock solution containing 5% (v/v) DMSO. For CD measurements, methanol was used as a co-solvent in place of DMSO to minimize solvent-associated spectral interference. The peptide solutions were incubated at room temperature overnight to allow supramolecular assembly prior to use. Stock solutions were diluted in Milli-Q water or cell culture medium as required for individual experiments. Working solutions were gently mixed immediately before use.

### Circular dichroism (CD) spectroscopy

CD spectra of peptide assemblies in aqueous solution were recorded at room temperature using a J-1700 CD spectrometer (JASCO, Japan) under a continuous nitrogen purge. Samples were loaded into a 1.0 mm path-length quartz cuvette (Starna, UK). Spectra were acquired from 185 to 350 nm at a scanning speed of 50 nm min⁻¹, with a digital integration time (D.I.T.) of 2 s, a data pitch of 0.5 nm, and a bandwidth of 1 nm. Baseline correction was performed using the corresponding solvent blank. Each spectrum represents the average of three consecutive scans. For presentation purposes, spectra were smoothed using a Savitzky–Golay filter (window length = 11 points, polynomial order = 3).

### Attenuated total reflection Fourier-transform infrared (ATR-FTIR) spectroscopy

ATR-FTIR spectra of lyophilized peptide assemblies were recorded using a Nicolet iS50 FTIR spectrometer (Thermo Fisher Scientific, USA). Samples were placed directly onto a diamond ATR crystal and spectra were collected at room temperature from 4000 to 400 cm⁻¹ with a spectral resolution of 1 cm⁻¹. Each spectrum represents the average of 64 scans. Atmospheric background correction was applied prior to analysis. Spectral processing was performed using OPUS 6.5 (Bruker, Germany), including sequential water vapor subtraction, baseline correction, and min–max normalization relative to the amide I region. The amide I band was analyzed by second-derivative deconvolution to identify component peaks. Peak fitting was performed using Gaussian functions, with peak widths constrained within comparable ranges across samples. Component peaks were assigned according to established amide I band positions corresponding to β-sheet, random coil, turn, and 3₁₀-helical structures.

### Zeta-potential measurement

Peptide solutions were incubated at room temperature overnight to allow supramolecular assembly prior to analysis. Samples were loaded into disposable folded capillary cells (DTS1070, Malvern, UK), and zeta potentials were measured in triplicate using a Zetasizer Nano ZS Red (Malvern, UK) by electrophoretic light scattering (ELS). Measurements were performed at room temperature with an equilibration time of 120 s.

### Rheology

Peptide solutions were prepared in Milli-Q water, pH-adjusted, and sonicated to obtain homogeneous dispersions. Samples were incubated at 37 °C for 48 h to induce self-assembly into hydrogels prior to measurements. Rheological characterization was performed using a parallel-plate geometry (100 μm gap) at 25 °C. Excess hydrogel was loaded onto the lower Peltier plate to ensure full coverage, and the upper plate was lowered to the measuring gap. Extruded material was trimmed to minimize edge effects and ensure consistent boundary conditions.

Strain amplitude sweeps were conducted at 6 rad s⁻¹ over 0.01–300% strain to determine the linear viscoelastic region (LVR). Frequency sweeps were then performed within the LVR over 100-0.01 rad s⁻¹ to obtain storage (G′) and loss (G″) moduli.

Stress relaxation measurements were performed by applying a constant strain selected from the overlapping LVR of all hydrogels and close to its upper limit. The strain was maintained for 180 s, and torque was recorded at 0.5 s intervals. Shear stress was calculated from torque using τ = 2M/(πR³) for parallel-plate geometry, and relaxation profiles were normalized to the initial value at 0.5 s. Relaxation dynamics were quantified by fitting the stress decay to a biexponential model based on a two-element Maxwell–Wiechert framework. The characteristic relaxation time (τ₁/₂), defined as the time required for stress to decay to 50% of its initial value, was extracted from fitted curves. Creep–recovery measurements were performed under a constant stress corresponding to 20% of the critical stress at the LVR boundary, followed by a 180 s recovery phase after stress removal.

### Transmission electron microscopy (TEM)

For TEM analysis, 5 μL of peptide solution was deposited onto plasma-treated carbon-coated copper grids (200 mesh; #D11022; KYKY Technology, China) and allowed to adsorb for 60 s. Excess solution was removed by gently blotting the grid edge with filter paper, followed by washing with Milli-Q water (3×) to remove unbound assemblies. The samples were then negatively stained with 1.0% (w/v) uranyl acetate for 10 s and air-dried at room temperature prior to imaging. The copper grids were pretreated using a COVANCE Vacuum Plasma System (Femto Science, Korea). TEM imaging was performed on a JEM-F200 field emission transmission electron microscope (JEOL, Japan) operated at 200 kV.

### Molecular dynamics (MD) simulations

Peptide structures were initially predicted using AlphaFold3 and subsequently converted into coarse-grained representations using the MARTINI 2.2 force field via Martinize2 (Vermouth framework). All residues were modeled in a random-coil state to avoid bias toward predicted secondary structures and to allow conformational flexibility during self-assembly. Each peptide carried a net charge of +1, corresponding to a protonated lysine residue. Simulations were performed using GROMACS 2025.2 in a cubic box containing 300 peptide molecules solvated with MARTINI water beads and 0.15 M NaCl, corresponding to a peptide concentration of ∼50 mM. Although higher than experimental conditions, this concentration is commonly used in coarse-grained simulations to enhance sampling of self-assembly processes.. After energy minimization, systems were equilibrated under NVT and NPT ensembles prior to production simulations. Production runs were performed in triplicate under NPT conditions at 303 K and 1 bar with periodic boundary conditions. Each trajectory was simulated for 2.5 μs. Molecular visualizations were generated using PyMOL.

### Cell culture

HeLa cells (RRID: CVCL_0030) were obtained from the Cell Bank of the Chinese Academy of Sciences (#TCHu187) and RIKEN BRC Cell Bank (#RCB0007) and cultured in DMEM supplemented with 10% (v/v) fetal bovine serum (FBS) and 1% penicillin/streptomycin. For experiments, cells in exponential growth phase were seeded into appropriate vessels and allowed to adhere for 8 h prior to treatment. All washes were performed using pre-warmed (37 °C) serum-free medium. For peptide treatment, stock solutions were diluted in low-serum medium (2% FBS unless otherwise stated) and applied to cells after thorough mixing. Equivalent volumes of sterile-filtered Milli-Q water served as controls.

### Cell viability assay

HeLa cells were seeded in 96-well plates at 1 × 10□cells per well and incubated with peptide assemblies at the indicated concentrations for 24–72 h. Cell viability was determined using the MTT assay. Briefly, MTT solution (5 mg mL⁻¹) was added and incubated for 4 h at 37 °C, followed by dissolution of formazan crystals in SDS solution. Absorbance was measured at 570 nm using a VICTOR Nivo Multimode Plate Reader (PerkinElmer, USA).

For long-term viability assessment, cells were seeded at reduced density and cultured for up to 7 days in the presence of peptide assemblies. Cell viability was quantified using a CCK-8 assay according to the manufacturer’s instructions, and absorbance was measured at 450 nm using a Synergy H1 Microplate Reader (BioTek, USA).

### Wound healing assay

Cell migration was evaluated using a wound-healing assay. Confluent HeLa monolayers were scratched using a standardized wound-making tool and subsequently treated with peptide assemblies at the indicated concentrations. Wound closure was monitored using an IncuCyte S3 live-cell imaging system (Essen BioScience, UK), and migration was quantified as relative wound density using the IncuCyte Scratch Wound Analysis module.

### Single-cell tracking assay

Cell motility was analyzed by time-lapse single-cell tracking. Following peptide treatment, nuclei were fluorescently labeled and imaged using a CellCyte live-cell imaging system at 30 min intervals for 24 h. Nuclear trajectories were reconstructed using TrackMate in Fiji, and migration distance and displacement were quantified. For Rho activation experiments, Rho Activator II was added following peptide treatment before imaging. For pyruvate rescue experiments, peptides were administered in culture medium supplemented with sodium pyruvate at either the normal (1 mM) or elevated (5 mM) concentration.

### Transwell invasion assay

Cell invasion was assessed using Matrigel-coated transwell inserts (8.0 μm pore size). Serum-starved HeLa cells were seeded into the upper chamber in the presence or absence of peptide assemblies, while complete medium containing 10% FBS was placed in the lower chamber as a chemoattractant. After 24 h, invaded cells were fixed, stained, and quantified by microscopy.

### Seahorse extracellular flux assay

Mitochondrial respiration and glycolytic activity were assessed using a Seahorse XF96 Extracellular Flux Analyzer (Agilent). HeLa cells were seeded in XF96 cell culture microplates and treated with 100 μM peptide as indicated. Oxygen consumption rate (OCR) and extracellular acidification rate (ECAR) were measured using the Seahorse XF Cell Mito Stress Test Kit (#103015-100) and Glycolysis Stress Test Kit (#103020-100), respectively, according to the manufacturer’s instructions.

For mitochondrial stress testing, cells were equilibrated in Seahorse XF assay medium supplemented with 10 mM glucose, 1 mM pyruvate, and 2 mM glutamine. Basal OCR was recorded followed by sequential injection of oligomycin (1.5 μM), FCCP (0.5 μM), and rotenone/antimycin A (0.5 μM). Basal respiration, ATP-linked respiration, maximal respiration, and spare respiratory capacity were calculated using Agilent Seahorse Analytics software. For glycolysis stress testing, cells were equilibrated in glucose-free Seahorse XF assay medium supplemented with 2 mM glutamine. ECAR was measured following sequential injection of glucose (10 mM), oligomycin (1 μM), and 2-deoxyglucose (50 mM).

### Glycolysis and oxidative phosphorylation (OXPHOS) assay

The relative contributions of glycolysis and oxidative phosphorylation (OXPHOS) to cellular energy production were evaluated using a Glycolysis/OXPHOS Assay Kit (Dojindo, #G270) according to the manufacturer’s instructions. HeLa cells were seeded in 96-well plates and treated with 100 μM **FFIKLLI** for 24 h under reduced-serum conditions (1% FBS). Cells were subsequently incubated with either oligomycin (1.25 μM) or 2-deoxyglucose (22.5 mM) for 5 h to selectively inhibit mitochondrial ATP synthesis or glycolysis, respectively. Lactate production was quantified from culture supernatants by absorbance measurement at 450 nm, while intracellular ATP levels were determined by luminescence. The relative contributions of glycolytic and mitochondrial metabolism were calculated according to the manufacturer’s protocol.

### Immunofluorescence staining (cell experiments)

Following the indicated treatments, HeLa cells cultured in glass-bottom confocal dishes were processed for immunofluorescence staining. Cells were fixed with 4% paraformaldehyde, quenched with ammonium chloride, permeabilized, and blocked before antibody incubation. Samples were incubated with primary antibodies overnight at 4 °C, followed by incubation with fluorophore-conjugated secondary antibodies for 1 h at room temperature. F-actin and nuclei were counterstained with ActinGreen 488 and DAPI, respectively. For Congo red staining, fixed cells were incubated with Congo red solution for 30 min before antibody staining.

To selectively detect cell surface–localized activated integrin β1, immunostaining with the HUTS-4 antibody was performed before permeabilization. Following incubation with fluorophore-conjugated secondary antibodies, cells were post-fixed, permeabilized, and subjected to intracellular immunostaining.

### Transfection and live-cell imaging

HeLa cells were transiently transfected with the indicated plasmids using Lipofectamine 3000 according to the manufacturer’s instructions. Transfected cells were subsequently reseeded into glass-bottom confocal dishes. Live-cell imaging was performed approximately 48 h after transfection using a stage-top incubation system maintained at 37 °C with 5% CO₂. Baseline images were acquired before addition of the FFIKLLI assemblies, and cells were imaged at the indicated time points thereafter. At the end of the experiment, nuclei were visualized using Hoechst 33342..

### Laser-scanning confocal microscopy and image analysis

Confocal images were acquired using a Nikon AX R laser-scanning confocal microscope mounted on an ECLIPSE Ti2-E inverted microscope equipped with 60× or 100× oil-immersion objectives. Images intended for quantitative comparison were acquired using identical imaging settings across experimental groups. Image denoising and deconvolution were applied for visualization only and were not used for quantitative analyses.

Cell morphology was quantified from F-actin-stained cells using Fiji. The ratio of phospho-myosin light chain (pMLC) to F-actin fluorescence intensity was calculated from maximum-intensity projections of z-stack images. The subcellular distribution of activated integrin β1 was analyzed from confocal z-stack images together with paxillin staining. The focal adhesion plane was defined as the z-position of maximal paxillin fluorescence intensity, and activated integrin β1 fluorescence was quantified relative to this plane. Fluorescence intensity was further quantified separately in the basal and apical plasma membrane compartments. Mitochondrial morphology was evaluated by TOM20 immunostaining using the Mitochondria Analyzer plugin in Fiji. Individual mitochondria were classified as filamentous, intermediate, or punctate according to established morphometric criteria, and the proportion of each category was determined. Mitochondria-associated DRP1 recruitment was quantified from TOM20/DRP1 co-immunostaining by measuring DRP1 fluorescence within TOM20-defined mitochondrial masks.

### Mitochondrial membrane potential assessment

Mitochondrial membrane potential was assessed using the JC-1 fluorescent probe. HeLa cells were treated with **FFIKLLI** for the indicated durations and subsequently stained with JC-1 and Hoechst 33342. Live-cell fluorescence imaging was performed under physiological culture conditions, and the relative abundance of JC-1 monomers and aggregates was quantified as an indicator of mitochondrial polarization state. Treatment with carbonyl cyanide *m*-chlorophenyl hydrazone (CCCP) served as a positive control for mitochondrial depolarization.

### Proteomic analysis

HeLa cells were cultured in 10-cm dishes to ∼50% confluency and subjected to the indicated treatments. Cells were then harvested and washed twice with ice-cold DPBS. Lysis was performed using buffer containing 20 mM HEPES, 8 M urea, and protease inhibitor cocktail (pH 8.0), followed by sonication on ice using a high-intensity ultrasonic processor (UR-21P; Tomy Digital Biology, Japan) with intermittent cooling to prevent overheating. Cell debris was removed by centrifugation (2.2 × 10□g, 10 min, 4 °C), and protein concentration was determined using a BCA assay (#23227; Thermo Scientific, USA). Quantitative proteomics was performed by PTM BIO (China). Briefly, proteins were digested with trypsin, and peptides were labeled using a TMT reagent kit (Thermo Scientific, USA) according to the manufacturer’s instructions. Labeled peptides were fractionated by high-pH reverse-phase HPLC using a ZORBAX 300Extend-C18 column (#770995-902; Agilent, USA). Peptide fractions were analyzed using an EASY-nLC 1000 system coupled to a Q Exactive Plus Orbitrap mass spectrometer (Thermo Scientific, USA) via a nanospray ionization source.

### Xenograft tumor model

All animal procedures were approved by the Animal Care and Use Committee of the Okinawa Institute of Science and Technology Graduate University. BALB/c nude (*nu/nu*) mice (female, 5 weeks old) were subcutaneously inoculated with HeLa-GFP cells (1 × 10 □ cells per mouse) to establish xenograft tumors. After tumor formation (∼50 mm³), animals were randomized into vehicle and **FFIKLLI**-treated groups (15 and 30 mg kg⁻¹, n = 5 per group). Peptide treatment was administered by local peritumoral injection once daily for 14 days under isoflurane anesthesia. Tumor progression was monitored longitudinally by caliper measurements and in vivo GFP fluorescence imaging (IVIS, PerkinElmer), enabling non-invasive tracking of tumor burden under identical acquisition parameters. Tumor volume was calculated as V = (L × W²)/2. At endpoint, tumors were harvested for weight measurement and histological processing following fixation in 4% paraformaldehyde.

### Hematoxylin and eosin (H&E) staining

Tumor tissues fixed in 4% paraformaldehyde were dehydrated through graded ethanol series and embedded in paraffin. Sections (5 μm) were prepared using a microtome (Leica, Germany) and mounted onto glass slides. After deparaffinization in xylene, sections were stained with hematoxylin for 5 min and eosin for 3 min, followed by rinsing in distilled water. Stained sections were imaged using a light microscope (Nikon, Japan).

### Immunofluorescence staining (tumor tissue sections)

For immunofluorescence analysis, deparaffinized sections were rehydrated in PBS and subjected to heat-induced antigen retrieval in 10 mM citrate buffer (pH 6.0). Sections were blocked with 5% BSA, followed by incubation with primary antibodies overnight at 4 °C. After washing, samples were incubated with fluorophore-conjugated secondary antibodies (1:1000) for 1 h at room temperature. Nuclei were counterstained with DAPI, and slides were mounted with antifade medium. Images were acquired using a confocal laser scanning microscope (A1 CLSM; Nikon, Japan).

### Statistical analysis and reproducibility

Measurements were obtained from one to three independent biological replicates unless otherwise stated. Representative images were selected from at least three independent experiments. Data are presented as mean ± s.d. Statistical analyses were performed using GraphPad Prism (v10.6.1) or the SciPy (scipy.stats) package in Python. The sample size (*n*) and statistical tests used are specified in the corresponding figure legends.

Comparisons between two groups were performed using two-tailed Student’s *t*-test or Welch’s *t*-test, depending on variance homogeneity. Multiple-group comparisons were performed using one-way Brown–Forsythe and Welch ANOVA followed by appropriate post hoc tests. Ternary compositional data were analyzed by MANOVA on centered log-ratio-transformed data. Significance thresholds are indicated in the corresponding figure legends.

## Acknowledgements

This work was supported by Songshan Lake Materials Laboratory, Guangdong S&T program (2024A0505040003), the National Natural Science Foundation of China (32500664), Guangdong Basic and Applied Basic Research Foundation (2025A1515010616), the Pearl River Talent Program of Guangdong Province (2023CX10C027), SLAB Young Scientists Program, SLAB AI+Materials Program. We thank Dr. Shijin Zhang and Dr. Sachie Yukawa at the Okinawa Institute of Science and Technology, Japan, for their assistance with the xenograft tumor studies and the H&E and immunohistochemical staining analyses.

## Author contributions

X.H. and Y.Z. conceived the study and designed the experiments. Q.Z., S.R., T.Z., W.H. and X.H. performed the experiments. Q.Z. conducted NMR, mass spectrometry, CD, and FTIR spectroscopy; prepared TEM and SEM samples; performed rheological characterization, EdU proliferation assays, glycolysis and OXPHOS analyses, transwell invasion assays, single-cell tracking assays, immunostaining and confocal microscopy, transfection and live-cell imaging, and mitochondria membrane potential and morphology analyses; S.R. conducted Zeta-potential measurements, wound healing assays, and Seahorse metabolic analyses. Q.Z. and S.R. performed cell viability assays, proteomic analyses, and Western blotting. T.Z. assisted with rheological data analysis and single-cell traction analysis. W.H. performed MD simulations. X.H. conducted MST measurements. C.X., J.Y. K.W. X.H., and Y.Z. advised the study. X.H. and Y.Z. wrote the maniscript with input from all authors.

## Competing interests

The authors declare the following competing financial interest(s): Q.Z., X.H. and Y.Z. are inventors on a patent application related to the data presented in this manuscript.

## Data and materials availability

All data are available in the main text or the supplementary materials.

