## Supplemental Text, table, figures for "Integrin-engaged Cellular Patches Mechanically Impose a Mitochondrial Respiratory Bottleneck to Suppress Cancer Cell Motility"

This PDF file includes:

Supplementary methods

Supporting figures S1-S30

Attachments 1-3

**Supplementary methods**

**Materials and Methods**

**Reagents**

Uranyl Acetate (depleted uranium; #02624-AB) from SPI Supplies (USA); Osmium Tetroxide (OsO_4_; #018456) from Ted Pella (USA); Sodium Cacodylate Trihydrate (≥ 98%; #205541) from Millipore (USA); Anhydrous Ethanol (EtOH; AR, ≥ 99.5%) from Nanjing Chemical Reagent (China); Methyl Alcohol (MeOH; HPLC, ≥ 99.90%; #75851G), Glutaraldehyde (25% in water; #14376J) and Triton X-100 (≥ 99%; #88963L) from Titan Scientific (China); Sodium Hydroxide (NaOH; ≥ 97.0%; #221465), Bovine Serum Albumin (BSA; ≥ 98%; #SRE0098) and Dimethyl Sulfoxide (DMSO; ≥ 99.7%; #D2650) from Sigma-Aldrich (USA); *tert*-Butanol (*t*-BuOH; AR, ≥ 99%; #A88563), Ammonium Chloride (NH_4_Cl; ≥ 99.5%; #A01043) and Dimethyl Sulfoxide-d6 (DMSO-*d*_6_; 99.8 atom % D; #D0125) from Innochem (China); 4% Paraformaldehyde (PFA) Fix Solution, QuickBlock Blocking Buffer (#P0260), QuickBlock Primary Antibody Dilution Buffer (#P0262) and SignalUp Secondary Antibody Dilution Buffer for Immunostaining (#P0278) from Beyotime (China). Unless otherwise specified, all reagents were used as received without further purification.

**Cell culture reagents** included Dulbecco's Modified Eagle Medium (DMEM, #11995-500), DMEM with low glucose (#11885-500), FluoroBrite DMEM Live Cell Fluorescence Imaging Medium (#A18967-01), OPTI-MEM I Reduced Serum Medium (#31985-062), TrypLE Express (#12605-010), Dulbecco's Phosphate Buffered Saline (DPBS; #14190-500), Penicillin-Streptomycin (P/S; 10,000 U mL^-1^; #15140-122), Sodium Pyruvate (100 mM; #11360-070) and GlutaMAX-I (100×; #35050-061) from GIBCO (USA); Fetal Bovine Serum (FBS; #F0193) from Sigma-Aldrich (USA); Insulin (#P3376) from Beyotime (China); Lipofectamine 3000 Transfection Kit (#L3000-008) from Invitrogen (USA); Rho Activator II (#CN03) from Cytoskeleton (USA); and Matrigel Growth Factor Reduced Basement Membrane Matrix (#354230) from Corning (USA).

**Fluorescent probes, staining reagents, and assay kits** included 3-(4,5-Dimethylthiazol-2-yl)-2,5-Diphenyltetrazolium Bromide (MTT; #M6494), NucBlue Live Cell Stain ReadyProbes reagent (Hoechst 33342, #R37605), NucBlue Fixed Cell Stain ReadyProbes reagent (DAPI, #R37606), ActinGreen 488 ReadyProbes reagent (#R37110), MitoProbe JC-1 Assay Kit (#M34152) from Invitrogen (USA); Crystal Violet Staining Solution (#C0121), Enhanced Cell Counting Kit-8 (#C0043), BeyoClick EdU Cell Proliferation Kit with AF488 (#C0071L) from Beyotime (China); PK Mito Deep Red (#PKMDR-2) from GenVivo Biotech (China); Glycolysis/OXPHOS Assay Kit (#G270) from Dojindo Laboratories (Japan); SPY595-DNA (#CY-SC301) from Spirochrome (Switzerland); and Congo Red amyloid fibril-binding dye (#ab145645) from Abcam (UK).

**Plasmids**

mEGFP-Lifeact-7 (RRID:Addgene_54610; Addgene plasmid #54610, http://n2t.net/addgene:54610) and DsRed2-Mito-7 (RRID:Addgene_55838; Addgene plasmid #55838, http://n2t.net/addgene:55838), both obtained as gifts from Michael Davidson.

**Primary antibodies**

Ms IgG_2a_ mAb to TOM20 (F-10, RRID:AB_628381; #sc-17764; 1:50) from Santa Cruz (USA); Rb IgG mAb to Paxillin (Y113, RRID:AB_779033; #ab32084; 1:50) and Rb IgG mAb to Drp1 (EPR19274, RRID:AB_2895215; #ab184247) from Abcam (UK); Rb pAb to Phospho-Myosin Light Chain 2 (pMLC, Thr18/Ser19, RRID:AB_2147464; #3674S; 1:200), Rb IgG mAb to Phospho-DRP1 (Ser616, D9A1, RRID:AB_11178659; #4494T) and Rb pAb to Phospho-DRP1 (Ser637, RRID:AB_10622027; #4867S) from Cell Signaling Technology (USA); and Ms IgG_2b_ mAb to Activated Integrin β1 (HUTS-4, RRID:AB_2233964; #MAB2079Z; 1:200) from Millipore (USA). **Secondary antibodies** used were: Goat pAb to Ms IgG (Alexa Fluor 647, RRID:AB_2687948; #ab150115; 1:1000), Dnk pAb to Ms IgG (Alexa Fluor 405, RRID:AB_2687445; #ab175658; 1:1000), Goat pAb to Rb IgG (Alexa Fluor 568, RRID:AB_2576207; #ab175471; 1:1000) and Dnk pAb to Rb IgG (Alexa Fluor 647, RRID:AB_2752244; #ab150075; 1:1000) from Abcam (UK).

**Instruments**

^1^H NMR and ^13^C NMR spectra were acquired on an Ascend Evo 500 Magnet System (500 MHz; Bruker, USA). The ^1^H NMR and ^13^C NMR chemical shifts $\delta$ are given in parts per million (ppm) referring to internal standard tetramethylsilane (TMS). Zeta-potentials were measured using a Zetasizer Nano ZS Red (Malvern, UK). Sample lyophilization was conducted using a YTLG-10C Desktop Lyophilizer (Shanghai Yetuo Technology, China). FTIR spectra were recorded on a Nicolet iS50 FTIR Spectrometer (Thermo Fisher Scientific, USA). CD were analyzed using a J-1700 CD Spectrometer (JASCO, Japan). Rheological measurements were performed using a Discovery Hybrid Rheometer (DHR) HR 30 (TA Instruments, USA) equipped with a Smart Swap 2 Stainless Steel Plate Geometry (20 mm diameter; #511200.946). TEM grids were pretreated in a COVANCE Vacuum Plasma System (Femto Science, Korea) and images were captured using a JEM-F200 TFEG Transmission Electron Microscope (JEOL, Japan). SEM samples were coated in an EM ACE600 Sputter Coater (Leica, Germany) and captured using a GeminiSEM 300 Field Emission Scanning Electron Microscope (Zeiss, Germany).

Cell lines were cultured in an MCO-230AICUVHL-PC CO_2_ Incubator (Panasonic, Japan). Colorimetric assays were performed using a VICTOR Nivo Multimode Plate Reader (PerkinElmer, USA) or a Synergy H1 Microplate Reader (BioTek, USA). Random cell migration was monitored by a Cellcyte X Live Cell Imaging System (Cytena, Germany). Fluorescent cell images were captured using an EVOS M7000 Imaging System (Invitrogen, USA). Confocal imaging was conducted by an AX R Confocal on an ECLIPSE Ti2-E Microscope (Nikon, Japan), using a 60×/1.42 (#MRD71670) or 100×/1.45 (#MRD71970) CFI Plan Apochromat Lambda D Oil Immersion Objective Lens, with a Bold Line Cage Incubator (Okolab, Italy).

**Rheology**

**Sample prepatation and rheometry setup**-Peptide samples were dissolved in Milli-Q water, pH neutralized, sonicated for a homogeneous dispersion and subsequently incubated at 37 °C for 48 h to promote self-assembly prior to rheological characterization. All measurements were conducted isothermally at 25 °C with a fixed gap of 100 μm. For each test, an excess amount of peptide hydrogel was loaded onto the lower Peltier plate to ensure full and uniform contact. The upper geometry was then gently lowered to the preset gap. Any material extruded beyond the plate edge was carefully trimmed with a laboratory spatula to obtain a consistent meniscus and to minimize extraneous viscous friction and torque contributions.

**Oscillatory rheology-**The viscoelastic behavior of the peptide hydrogels was characterized using a multi-step protocol. Oscillatory **strain amplitude sweeps** were first conducted at a fixed angular frequency of 6 rad/s over a strain range of 0.01%–300% with 10 logarithmically spaced data points per decade, to establish the linear viscoelastic region (LVR). Oscillatory **frequency sweeps** were then performed within the LVR across 100–0.01 rad/s with 10 logarithmically spaced data points per decade, to obtain the storage modulus $G'$ and the loss modulus $G''$.

**Stress relaxation and analysis**-The tests under shear deformation were performed by applying an identical strain selected from the overlapping LVR of all peptide hydrogels and close to its upper limit to maximize signal sensitivity while maintaining linear viscoelastic conditions. The strain was held constant and the decay of shear torque was recorded at 0.5 s intervals over 180 s to evaluate viscoelastic relaxation behavior. As the shear stress $\tau$ is directly proportional to torque $M$ for a parallel-plate geometry with known radius $R$ as $\tau=\frac{2M}{\pi R^{3}}$, normalized stress relaxation profiles were represented by torque normalized to the first recorded data point (0.5 s).

To find $\tau_{1/2}$ (the time that gels took to relax their initial stress to its half value) of different gels, stress-relaxation curves were fitted into a biexponential equation based on a two-element Maxwell-Weichert model and $\tau_{1/2}$ was calculated from each curve.

**Creep-recovery**-The tests consisted of two consecutive steps^1^. In the creep phase, a constant shear stress was applied for 180 s. For each sample, the applied stress was selected as 20% of the critical stress corresponding to upper boundary of LVR, as determined from strain sweep measurement, to ensure linear viscoelastic conditions while maintaining sufficient deformation sensitivity. Subsequently, the applied stress was removed, and the recovery phase continued for an additional 180 s. Deformation was recorded as rotational displacement $\theta$ under the fast sampling mode throughout the whole test, from which shear strain $\gamma$ was calculated for the parallel-plate geometry according to $\gamma=\frac{\theta R}{h}$, where $R$ and $h$ represent the plate radius and gap, respectively. Creep compliance $J(t)$ and recoverable compliance $J_{r}(t)$ were calculated from the creep strain $\gamma(t)$ and the recovered strain $\gamma_{r}\left( t \right)$, respectively, with $\gamma_{r}\left( t \right)=\gamma_{max}-\gamma(t)$ during the recovery phase, and normalized by the applied shear stress $\sigma$ during the creep phase, yielding $J\left( t \right)=\frac{\gamma(t)}{\sigma}$ and $J_{r}\left( t \right)=\frac{\gamma_{max}-\gamma(t)}{\sigma}$.

**Transmission electron microscopy (TEM)**

To improve the readability of several TEM micrographs containing locally electron-dense regions arising from fiber over-accumulation, a Gaussian blur–based background subtraction was applied in Fiji^2^. Briefly, a duplicate of each image was blurred with a Gaussian filter ($\sigma=100 \mathrm{pixels}$), and the resulting background estimate was subtracted from the original image using the Image Calculator function.

**Molecular dynamics (MD) simulation**

Peptide structures were first predicted using AlphaFold3^3^ and subsequently converted into a MARTINI 2.2 coarse-grained (CG) model^4^ via Martinize2 within the Vermouth framework^5^. To avoid introducing secondary structure bias into the self-assembly simulations and allow for conformational flexibility, all residues were assigned as random coil during CG mapping, regardless of the helical conformations predicted by AlphaFold3. All three peptide sequences (**IKLLI**, **FFIKLLI**, and **FFILIKL**) contain only one charged residue, the naturally protonated lysine (K, +1). No protonation state modifications were applied during coarse-graining, and the default protonation states defined in the MARTINI 2.2 force field were retained. As a result, each peptide carried a net charge of +1.

Simulations were performed in a cubic box of 21.5 × 21.5 × 21.5 nm containing 300 peptide molecules, corresponding to a peptide concentration of approximately 50 mM. Although this exceeds typical biological experimental conditions at concentrations on the order of 100 μM, such elevated concentrations are well established in CG-MD simulations of peptide self-assembly to enhance sampling efficiency within accessible simulation timescales^6^. Systems were solvated with standard MARTINI water beads and supplemented with 0.15 M NaCl to mimic physiological ionic strength and maintain overall charge neutrality.

Energy minimization was carried out using the steepest descent algorithm for up to 5000 steps, or until the maximum force fell below 1000 kJ mol^-1^ nm^-1^, with positional restraints of 2000 kJ mol^-1^ nm^-2^ applied to backbone beads. A cutoff of 1.1 nm was used for all nonbonded interactions. Electrostatic interactions were treated using the reaction-field method with a relative dielectric constant of 15. The Lennard–Jones potential was handled with a potential-shift scheme as defined in the MARTINI 2.2 force field^7^.

Following energy minimization, the system was equilibrated in two consecutive phases to gently relax the solvent around the solute without inducing structural distortion. First, a 1 ns isochoric-isothermal (NVT) equilibration was performed using a 20-fs integration time step. Initial velocities were generated according to a Maxwell–Boltzmann distribution at 303 K. The temperature was maintained at 303 K using the V-rescale thermostat with a coupling time constant of 1.0 ps. Subsequently, a 10 ns isobaric-isothermal (NPT) equilibration was conducted. To avoid drastic volume oscillations during initial pressure coupling, the pressure was isotropically coupled to 1.0 bar using the first-order C-rescale barostat with a coupling time of 12.0 ps and an isothermal compressibility of 3 × 10^-4^ bar^-1^. Throughout both the NVT and NPT equilibration phases, positional restraints of 1000 kJ mol^-1^ nm^-2^ were continuously applied to the peptide backbone beads.

Subsequent production CG-MD simulations were performed using GROMACS (v2025.2)^8^ in the NPT ensemble with a 25-fs integration time step and without any positional restraints. Temperature was maintained at 303 K using the V-rescale thermostat ($\tau_{T}=1 ps$)^7^, and pressure was controlled at 1 bar using the Parrinello–Rahman barostat ($\tau_{P}=12 \mathrm{ps}$) to yield correct thermodynamic ensemble fluctuations, with an isothermal compressibility of 4.5 × 10^-5^ bar^-1^. Periodic boundary conditions were applied in all three directions for all simulations. Each system was simulated for 10^8^ steps, corresponding to a 2.5-μs CG-MD production run^9^. Each simulation was performed in triplicate using different initial random seeds for Maxwell–Boltzmann velocity generation, effectively serving as independent production runs to mimic experimental replicates and ensure statistical reproducibility. All molecular visualizations were generated with PyMOL 2.5 (Schrödinger, USA).

**MD simulation analysis**

To evaluate the self-assembly behavior and structural stability of peptide assemblies, four categories of post-simulation analyses were performed on the CG-MD trajectories of all three sequences (**IKLLI**, **FFIKLLI**, and **FFILIKL**), except where noted. Time-resolved analyses were performed using trajectories sampled every 10^4^ simulation steps from the 10^8^-step production runs.

**Clustering analysis.** The temporal evolution of aggregate morphology was monitored by tracking two clustering metrics throughout the entire simulation trajectory: the maximum cluster size, defined as the number of peptide chains contained in the largest cluster at each time point, and the total number of clusters. Cluster analysis was performed using *gmx clustsize* with a distance cutoff of 0.5 nm. Both metrics were computed for all three sequences, including **IKLLI**, for which the analysis served as a baseline to confirm the absence of stable aggregate formation under the simulation conditions.

**Energy analysis.** The time evolution of four energy components, namely bond stretching energy (Bond), angle bending energy (G96Angle), short-range Lennard–Jones interaction energy (LJ_SR), and short-range Coulomb interaction energy (Coulomb_SR), together with the total potential energy (Total Potential) and total energy (Total Energy), was monitored throughout the simulation trajectory for all three sequences. Bond stretching energy reflects constraints imposed by covalent connectivity, whereas angle bending energy characterizes local conformational preferences. Non-bonded interactions were evaluated through short-range Lennard–Jones interaction energy, representing hydrophobic and steric contributions, and short-range Coulomb interaction energy, representing electrostatic interactions. Total potential energy and total energy were used to assess overall thermodynamic stability and simulation convergence.

**Per-residue RMSF.** Root mean square fluctuation (RMSF) was calculated on a per-residue basis for **FFIKLLI** and **FFILIKL** using the final one-fifth of each trajectory, corresponding to the equilibrated regime of the simulation. Prior to RMSF calculation, trajectories were centered and rotationally aligned to the peptide aggregate reference structure to remove contributions from global translational and rotational diffusion of the assembly. RMSF values were averaged across all peptide chains within the aggregate. RMSF analysis was not performed for **IKLLI** because no stable aggregate was formed under the simulation conditions.

**Per-residue SASA.** Solvent-accessible surface area (SASA) was computed on a per-residue basis for **FFIKLLI** and **FFILIKL** over the final one-fifth of each trajectory using *gmx sasa* with a custom van der Waals radii file adapted for MARTINI 2.2 CG beads, in which regular and small beads were assigned radii of 0.264 nm and 0.230 nm, respectively. Values were averaged across all peptide chains. SASA analysis was not performed for **IKLLI** because no stable aggregate was formed under the simulation conditions.

**Scanning electron microscopy (SEM)**

For the preparation of the SEM samples, 4 × 10^3^ HeLa cells were seeded in the 14 mm glass bottom well of a 35 mm cell culture dish (#D11130H; Matsunami Glass, Japan). After 24 h incubation with peptide assemblies, the culture medium was removed, and the cell culture was washed once. Pre-warmed 2.5% glutaraldehyde in 0.1 M cacodylate buffer (pH 7.4) was added to the cells for 10 min to fix the cells, followed by washing with 0.1 M cacodylate buffer thrice for 5 min each. The cells were post-fixed with 1% OsO_4_ (w/v) in 0.1 M cacodylate buffer for 30 min to stabilize membrane lipids, followed by washing with double distilled water (ddH_2_O) thrice for 5 min each. The samples were then dehydrated through a graded ethanol series (70%, 80%, 90%, 95% and 100%; 3 changes of 5 min each, with 100% repeated for 3 rounds), and displaced into *t*-BuOH thrice for 5 min each. Approximately 30% of *t*-BuOH was left at the last step to avoid cracking the biosamples, which were subsequently froze at -80 °C and lyophilized overnight. Glass coverslips retrieved from the dishes were mounted onto copper stubs using double-sided adhesive carbon tape (#05072-AB; SPI Supplies, USA) and sputter-coated with 3 nm of platinum (Pt), before SEM imaging at 1.0 kV with a working distance of approximately 7 mm.

**Cell viability assay**

HeLa cells were seeded in 96-well plates at a density of 1 × 10⁴ cells per well. After attachment, culture medium was replaced with medium containing peptide assemblies at final concentrations of 5, 10, 25, 50, or 100 μM. Cells were incubated for 24, 48, or 72 h.

For MTT assays, 10 μL of MTT solution (5 mg mL⁻¹) was added to each well and incubated for 4 h at 37 °C. The reaction was terminated by addition of 100 μL of 10% (w/v) SDS solution, followed by overnight incubation to dissolve formazan crystals. Absorbance was measured at 570 nm using a VICTOR Nivo Multimode Plate Reader (PerkinElmer, USA).

For long-term viability studies, 1250 or 750 cells per well were seeded for 5- or 7-day assessments, respectively. At the indicated time points, CCK-8 reagent was added and allowed to react for 1.5 h. Eighty microliters of supernatant was transferred to a new plate and absorbance was measured at 450 nm using a Synergy H1 Microplate Reader (BioTek, USA). Data are presented as mean ± s.d. from at least three independent experiments.

**Wound healing assay**

HeLa cells were seeded in 96-well plates at 3.5 × 10⁴ cells per well and cultured to approximately 90% confluence. Cells were serum-starved overnight in medium containing 2% FBS. Uniform scratch wounds were generated using an IncuCyte 96-Well WoundMaker Tool (Essen BioScience, UK). Detached cells were removed by washing three times with culture medium.

Cells were treated with peptide assemblies at concentrations of 5–100 μM. Images were acquired every 12 h for 60 h using an IncuCyte S3 live-cell imaging system (Essen BioScience, UK). Wound closure was quantified as relative wound density using the IncuCyte Scratch Wound Analysis module.

**Single-cell tracking assay**

Cells seeded in 96-well plates at a density of 2 × 10^3^ cells per well were incubated with or without treatment of 100 µM **FFIKLLI** for 24 h. SPY595-DNA was added to label the cells 2 h prior to the end of the 24 h treatment. Following treatment, cells were imaged using a 10× objective on a Cellcyte live cell imaging system. Time-lapse imaging was conducted for 24 h at 30 min intervals. After performing slice registration by Template Matching plugin^10^ in Fiji, the centroids of fluorescently labeled nuclei were tracked using the TrackMate plugin^11^ in Fiji, obtaining coordinates at each time point, total distance travelled, and total displacement for each cell. Cells with incomplete tracking records were excluded from further analysis. A custom Python script was used to reconstruct cell migration trajectories by sequentially linking the coordinates across consecutive time points for each cell, and to visualize all trajectories on a polar coordinate grid using the *matplotlib.pyplot* library (v3.1.1)^12^. For Rho activation assay, after the 24 h peptide treatment, SPY595-DNA along with or without Rho Activator II at final concentration of 1 µg mL^-1^ were added to the cells. Time-lapse imaging was initialized 2 h later. For pyruvate rescue assay, FFIKLLI were diluted in culture medium supplemented with sodium pyruvate at normal (1 mM) or high (5 mM) concentration for cell treatment.

**Transwell invasion assay**

Cell invasion was assessed using Matrigel-coated transwell inserts with 8.0 μm pore polycarbonate membranes (Corning, USA). Inserts were coated with 100 μL Matrigel (250 μg mL⁻¹) and allowed to gel at 37 °C for 2 h.

HeLa cells were serum-starved in medium containing 2% FBS for 12 h before collection. Cells (4 × 10⁴) suspended in serum-free medium containing the indicated peptide assemblies were added to the upper chamber. Complete medium containing 10% FBS was added to the lower chamber as a chemoattractant.

After 24 h, invaded cells were fixed in methanol, stained with crystal violet, and imaged using an ECLIPSE Ts2R-FL microscope (Nikon, Japan). DAPI staining was subsequently performed, and invaded cells were quantified using Fiji.

**Seahorse extracellular flux analysis**

Oxygen concentration rate (OCR) and extracellular acidification rate (ECAR) were measured using a Seahorse XF Extracellular Flux Analyzer (Agilent, USA). One day prior to the assay, the analyzer was set at 37 °C temperature overnight for equilibration. Meanwhile, Seahorse XF96 biosensor cartridge was hydrated with 180 µL of Seahorse XF calibration buffer (#100840) and incubated overnight at 37 °C with non-CO_2_ environment. On the day of the assay, the hydrated cartridge containing calibrant buffer was loaded into the analyzer for sensor calibration prior to the start of the experiment. HeLa cells were seeded at a density of 10^5^ cells mL^-1^ in 175 μL per well in Seahorse XF96 Cell Culture Microplates and treated with 100 μM peptide solution of the same volume for 12 h for mitochondrial respiration assay and in a time-dependent manner for glycolytic function assay.

**Mitochondrial respiration** was assessed using the Agilent Seahorse XF Cell Mito Stress Test Kit (#103015-100). The cell culture medium was replaced with 180 μL pre-warmed (37 °C) Seahorse XF Assay Medium (pH 7.4; #103575-100) supplemented with 10 mM Seahorse XF Glucose (#103577-100), 1 mM Seahorse XF Pyruvate (#103578-100), and 2 mM Seahorse XF Glutamine (#103579-100), and cells were equilibrated for approximately 1 h at 37 °C in a non-CO_2_ incubator prior to measurement. Basal OCR (pmol/min) was recorded over three measurement cycles, followed by sequential injections of oligomycin (final well concentration of 1.5 µM), carbonyl cyanide-*p*-trifluoromethoxyphenylhydrazone (FCCP; 0.5 µM), and rotenone/antimycin A (Rot/AA; 0.5 µM) at 17.10, 37.25 and 57.43 min, respectively. The key parameters of mitochondrial function were calculated automatically using Agilent Seahorse Analytics software, including basal respiration (baseline OCR − non-mitochondrial OCR), ATP production (basal OCR − oligomycin-treated OCR), maximal respiration (FCCP-stimulated OCR − non-mitochondrial OCR) and spare respiratory capacity (maximal respiration − basal respiration).

**Glycolytic function** was assessed using the Seahorse XF Glycolysis Stress Test Kit (#103020-100). Cells were equilibrated in 175 μL glucose-free XF assay medium supplemented with 2 mM glutamine for approximately 1 h at 37°C in a non-CO_2_ incubator prior to the assay to ensure glucose-starved baseline conditions. OCR and ECAR (mpH/min) were measured with sequentially injections of glucose (final well concentration of 10 mM), oligomycin(1 µM), and 2-deoxyglucose (2-DG; 50 mM) at the same time points as mitochondrial respiration assay.

**Glycolysis/OXPHOS assay**

Cellular ATP level and lactate production were evaluated using a glycolysis/OXPHOS assay kit (#G270; Dojindo, Japan) as per the manufacturer's instructions. In brief, 5 × 10^3^ cells were seeded into a 96-well white microplate (#3917; Coning, USA) and treated with or without 100 µM **FFIKLLI** in culture medium with 1% FBS for 24 h. 10× mitochondrial OXPHOS inhibitor oligomycin or glycolysis inhibitor 2-DG with final concentrations of 1.25 µM or 22.5 mM, respectively, was added to the culture medium and cells were stimulated for further 5 h. Thereafter, 20 μL of supernatant was diluted 10-fold in Milli-Q water, and 20 μL of each diluted sample was transferred to a clear microplate, followed by the addition of 80 μL lactate working solution using a multichannel pipette. The reaction mixture was incubated at 37 °C for 30 min and the absorbance at 450 nm was measured using a Synergy H1 Microplate Reader (BioTek, USA). For ATP level measurements, to the remaining medium in the white microplate, 100 μL ATP working solution was added. The plate was shake for 2 min and allowed to stand at 25 °C in for 10 min, before luminescence measurement using the same microplate reader. Data are presented as relative cellular ATP levels and lactate production, normalized to the untreated, inhibitor-free control group.

**Immunofluorescence (IF) staining (cell experiments)**

4 × 10^3^ HeLa cells in 200 μL culture medium were seeded in the 15 mm glass bottom well of a 35 mm cell culture dish (confocal dish; #801002; NEST Biotechnology, China) and treated accordingly. After removal of culture medium and rinse, cells were fixed with pre-warmed (37 °C) 4% PFA for 15 min at RT and washed twice with DPBS, followed by incubation with 0.1 M NH_4_Cl in DPBS for 10 min at RT to quench residual aldehyde group and washing twice with DPBS. Permeabilization and blocking of nonspecific binding sites were performed with QuickBlock blocking buffer for 20 min at RT. The cells were incubated with primary antibodies in QuickBlock primary antibody dilution buffer for 14 h at 4 °C. The unbound antibodies were washed away with 0.1 % PBS-T thrice at 5 min each. Cells were then incubated with secondary antibodies prepared in SignalUp secondary antibody dilution buffer for 1 h at RT in the dark. After washing thrice for 5 min each with 0.1% PBS-T, ActinGreen 488 and DAPI diluted in 0.1% PBS-T were used to stain the F-actin and nuclei respectively, for 20 min at RT in the dark. Cells were finally washed thrice with 0.1% PBS-T and replaced with 750 μL PBS and ready for imaging. For experiments involving intensity quantification, imaging was performed within 24 h of staining.

For Congo red staining of peptides, fixed cells were firstly incubated with 0.1 mg mL^-1^ freshly prepared Congo red in DPBS for 30 min at 37 °C. Cells were washed thrice with DPBS prior to further staining.

To restrict integrin staining to the plasma membrane, IF staining for activated integrin β1 was performed prior to permeabilization. Briefly, fixed cells were blocked with 3% (w/v) BSA in DPBS for 45 min at RT, then incubated with anti-activated integrin β1 antibody (HUTS-4; 1:200) in DPBS containing 1% BSA for 1 h at 4 °C. After three washes with DPBS, cells were incubated with goat anti-mouse IgG secondary antibody (Alexa Fluor 647; 1:1000) in 1% BSA for 1 h at RT in the dark. Following three washes with DPBS, cells were post-fixed with 4% PFA for 10 min at RT to immobilize the antibody complex, washed twice with DPBS, and incubated with 0.1 M NH_4_Cl in DPBS for 10 min at RT. Cells were then washed twice with DPBS before proceeding to permeabilization and further IF staining for intracellular proteins.

**Transfection and live-cell imaging**

For transient transfection, HeLa cells were seeded in a 6-well plate at a density of 5 × 10^5^ cells per well and allowed to adhere overnight. Prior to transfection, cells were washed twice with serum- and antibiotics-free medium and maintained in 1.25 mL of the same medium. Meanwhile, 2.5 μg plasmid DNA was mixed well with 3.75 μL Lipofectamine 3000 reagent in a total of 250 μL OPTI-MEM medium and incubated for 15 min at RT before added to the cells. Cells were incubated with the transfection solution for 6 h, after which they were washed twice with serum- and antibiotics-free medium and further incubated in antibiotics-free medium supplemented with 10% FBS.

At 36 h post-transfection, cells were reseeded into confocal dishes at a density of 1 × 10^4^ cells per well. At 44 h post-transfection, the medium was replaced with FluoroBrite DMEM supplemented 2% FBS, 4 mM GlutaMAX, and 1 mM sodium pyruvate for equilibration. Cells were imaged about 48 h post-transfection at 37 °C under 5% CO_2_ in a fully humidified Bold Line Cage Incubator (Okolab, Italy). Fields of interest were selected and *t* = 0 images were acquired, after which **FFIKLLI** was added at a final concentration of 100 μM and cells were imaged at 2, 6, 12, 18, and 24 h post-treatment. Following the final time point, cells were stained with Hoechst 33342 for nuclear visualization.

**Laser-scanning confocal microscopy and image analysis**

Confocal imaging was performed on an AX R Confocal system mounted on an ECLIPSE Ti2-E inverted microscope (Nikon, Japan), equipped with a LUA-S4 Laser Unit (405/488/561/640 nm; #MHF490AA) and PMT-GaAsP Detector Units (#MHE60220). Simultaneous differential interference contrast (DIC) imaging was acquired using an AX-DUT Diascopic Detector Unit (#MHE50350). Images were acquired using a 60×/1.42 NA (#MRD71670) or 100×/1.45 NA (#MRD71970) CFI Plan Apochromat Lambda D Oil Immersion Objective Lens with non-drying immersion oil for fluorescence microscopy (Type HF; #16245; Cargille Laboratories, USA). All imaging was performed through #1.5 glass (0.17 mm). Images were acquired in Galvano scanning mode with a pixel dwell time of ≥ 1 μs, ≥ 2-frame averaging, and a pinhole of ≤ 1 AU (referenced to the longest-wavelength channel) to improve signal-to-noise ratio. Channels were imaged sequentially from longest to shortest wavelength to minimize crosstalk. For experiments involving fluorescence intensity quantification, identical acquisition settings were applied across all groups, including laser power, detector gain, pinhole size, and sampling density (image resolution and zoom factor). For channels not intended for fluorescence intensity quantification, the *Restore.ai* denoising and deconvolution function in NIS-Elements ER software (v6.02.03) was applied for visualization purposes only.

For cell morphological analysis, a single optical slice from the ventral stress fiber–rich plane was acquired from each F-actin–stained cell. Cell contours were delineated using the Versatile Wand Tool plugin in Fiji with value tolerance of 100 to identify the background region, and morphometric measurements were performed with the resulting cell boundary.

For quantification of the phospho-myosin light chain (pMLC)–to–F-actin fluorescence intensity ratio, cells were immunostained with anti-pMLC antibody (Thr18/Ser19; 1:200), followed by F-actin labeling with ActinGreen 488. Maximum intensity projection (MIP) images were generated from five consecutive *z*-slices acquired at 0.5 μm intervals centered on the ventral stress fiber–rich plane. Fluorescence intensity analysis was performed on MIP images, restricted within cell boundary derived from the F-actin channel.

For localization of activated integrin β1, cells were immunostained with anti-activated integrin β1 (HUTS-4; 1:200) and anti-paxillin (1:50) antibodies, followed by nuclear and F-actin labeling with DAPI and ActinGreen 488, respectively. Consecutive z-slices of F-actin, paxillin, and integrin β1 channels were acquired at 0.48 μm intervals from the apical to the basal plasma membrane. The focal adhesion plane was defined as the *z*-slice containing maximal paxillin intensity and assigned as *z* = 0, with all relative *z*-positions expressed in μm with respect to this reference. Profiles of activated integrin β1 immunofluorescence along the *z*-axis were presented. Skewness was calculated as the normalized third central moment of the fluorescence intensity–weighted *z*-position distribution. The basal plasma membrane region was defined as the *z*-slices spanning the focal adhesion plane and extending up to three slices above it; whereas *z*-slices beyond this boundary were classified as the apical plasma membrane region. Fluorescence intensities of activated integrin β1 were quantified from MIP images of the whole cell as well as from each of the two defined compartments.

For mitochondrial morphometric analysis, the mitochondrial outer membrane protein TOM20 was immunostained with anti-TOM20 antibody (1:50), followed by nuclear and F-actin labeling with DAPI and ActinGreen 488, respectively. The scan area was adjusted to accommodate a full cell, with rotation applied where necessary to achieve a zoom factor of ~2.1× for Nyquist sampling criterion. The TOM20 channel was acquired at the z-slice where mitochondria in the periphery of cells were clearly resolved, using a pixel dwell time of 8 μs and 4-frame averaging, followed by deconvolution. Images were then contrast-enhanced (saturated pixels = 0.35%), converted to 8-bit, and background noise removed by setting pixels below an intensity of 30 to zero. A binary mask derived from the F-actin channel was subsequently applied to the TOM20 channel to restrict the signal to a single cell. The preprocessed images were then submitted to Mitochondria Analyzer plugin^1^ in Fiji for mitochondrial segmentation. Briefly, the TOM20 channel was binarized using the *2D Threshold* command with the Weighted Mean method (*block size* = 1.85 μm; *C-value* = 11), with all pre- and post-processing parameters set to default. Morphometric quantification was performed using the *2D Analysis* command at both per-cell and per-mitochondrion basis. For each cell, mitochondria were classified as filamentous (branches ≥ 3), punctate (branches = 1 and aspect ratio ≤ 2), or intermediate. The area fraction occupied by each category was calculated and presented as stacked bar plots and ternary plots.

For analysis of mitochondrial dynamin-related protein 1 (DRP1), TOM20 and DRP1 were simultaneously immunostained with anti-TOM20 and anti-DRP1 (1:250) antibodies, followed by F-actin labeling with ActinGreen 488. Mitochondrially associated DRP1 was isolated by generating a binary mask from the TOM20 channel and applying it to exclude all extra-mitochondrial DRP1 fluorescence signal.

**Mitochondrial membrane potential assessment**

Mitochondrial membrane potential was assessed using the JC-1 dye (5,5’,6,6’-tetrachloro-1,1’,3,3’-tetraethyl benzimidazolylcarbocyanine iodide). 2 × 10^4^ HeLa cells were seeded in confocal dishes and treated with or without 100 μM **FFIKLLI** for 24 or 48 h. Cells were then incubated with JC-1 (5 μg mL^-1^) and Hoechst 33342 in culture medium supplemented with 2% FBS for 30 min. After two washes, cells were imaged in FluoroBrite DMEM at 37 °C under 5% CO_2_. Fluorescence was acquired at excitation/emission of 488/525 ± 26 nm for JC-1 monomers and 561/598 ± 27 nm for JC-1 aggregates, to distinguish polarized from depolarized mitochondrial states. Multi-cell images were captured. Fluorescence intensity analysis was restricted within cell regions delineated by selection of the background in the JC-1 monomer channel. For mitochondrial depolarization control, JC-1-stained untreated group was incubated with 10 μM CCCP (carbonyl cyanide *m*-chlorophenyl hydrazone) in FluoroBrite DMEM for 20 min before immediate imaging.

**Bioinformatics analysis**

The resulting MS/MS data were processed using Maxquant search engine (v1.5.2.8)^14^. Tandem mass spectra were searched against the Swiss-Prot Human database^15^ concatenated with reverse decoy database. The mass tolerance for precursor ions was set as 20 ppm in *First search* and 5 ppm in *Main search*, and the mass tolerance for fragment ions was set as 0.02 Da. Identified proteins with the UniProt accession numbers were assigned to Gene Ontology (GO) terms^16^ according to the UniProt-GOA database^17^. If some identified proteins were not annotated by UniProt-GOA database, the InterProScan^18,19^ would be used to annotated protein’s GO functional based on protein sequence alignment method.

**Clustering analysis.** Proteins with statistically significant changes (*P* < 0.05) in relative abundance across at least one time point compared to the control group were selected for hierarchical clustering analysis. The filtered relative abundance ratios were subjected to unsupervised hierarchical clustering using Euclidean distance and complete linkage, and visualized as a heatmap using the *heatmap.2* function from the *gplots* package in R (v3.5.0). Row ordering was determined by the clustering result, and proteins were subsequently annotated into functional categories (cytoskeleton-, integrin-, and metabolic-related) based on their cluster assignment and known biological functions.

**GO enrichment analysis.**

The proteins were classified by GO annotation^20^ based on three categories: biological process, cellular component, and molecular function. For each category, a two-tailed Fisher’s exact test was employed to test the enrichment of the differentially expressed protein against all identified proteins. The GO with a corrected *P* < 0.05 value is considered statistically significant.

**Subcellular localization** of the identified proteins was predicted using the WoLF PSORT software^21^. Based on their amino acid sequences, proteins were assigned to distinct cellular compartments such as the extracellular space, plasma membrane, cytoplasm, nucleus, mitochondria, endoplasmic reticulum (ER), Golgi apparatus, peroxisome, lysosome, and cytoskeleton, with dual localizations (e.g., cyto_nucl, mito_pero) designated for proteins predicted to occupy multiple compartments.

**Xenograft tumor model**

All experimental procedures were performed in accordance with and approved by the Okinawa Institute of Science and Technology Graduate University Animal Care and Use Committee. 4-week-old female BALB/c nude (*nu*/*nu*) mice were purchased from Shizuoka Laboratory Center (SLC; Japan) and fed for another week before tumor development. HeLa-GFP cells in exponential growth phase were collected and suspended in 3 × 10^7^ cells/mL suspension. 100 µL of HeLa-GFP suspension was subcutaneously injected into the right flank of each nude mouse.

After 2 weeks of inoculation, mice bearing HeLa-GFP xenograft tumors with a mean tumor volume of approximately 50 mm^3^ were randomly assigned into a vehicle control group and two treatment groups, with 5 mice per group. Subcutaneous peritumoral injections of saline, 15, or 30 mg/kg **FFIKLLI** were administered once daily for 14 consecutive days under 2% isoflurane anesthesia with the first injection designated as *Day* *0*. Body weight and tumor dimensions were recorded before each injection. Tumor volume $V$ was calculated by caliper measurements using the formula $V=\frac{L\times W^{2}}{2}$, where $L$ and $W$ denote the longest and shortest diameters of the tumor, respectively. Tumor progression was additionally assessed by fluorescence imaging on *Days* *0*, *7*, and *14*, where anesthetized nude mice bearing HeLa-GFP tumors were imaged using a Lumina K In Vivo Imaging System (IVIS; PerkinElmer, USA) with a GFP fluorescence filter set and identical imaging parameters applied across all time points.

On *Day 14*, following the final routine assessments, mice were sacrificed and tumors were carefully excised, washed with ice-cold saline, weighed, and photographed alongside a ruler for size reference. Dissected tumor tissues were then fixed in 4% paraformaldehyde for 48 h at 4 °C prior to further processing.

**Statistical analysis and reproducibility**

No statistical methods were used to predetermine sample size. The experiments were not randomized, and the investigators were not blinded to group assignment during data collection or analysis. All measurements were performed on 1−3 independent biological replicates from separate experiments. All representative micrographs were selected from data sets from at least three independent experiments. Statistical analyses were carried out using GraphPad Prism (v10.6.1; GraphPad Software, USA) or *scipy.stats* library (v1.5.2)^2^ in Python. Data are presented as mean ± standard deviation (s.d.), as indicated in the figure legends. The exact sample size (*n*) and statistical test used for each data set are specified in the corresponding figure legends.

For the comparison of two unpaired data sets, homoscedastic data ($\frac{1}{2}⁠\leq⁠\frac{s_{1}}{s_{2}}⁠\leq2$) were analyzed by two-tailed Student’s *t*-test, whereas heteroscedastic data were analyzed by two-tailed Welch’s *t*-test. To compare more than two data sets with one factor, homoscedasticity was inspected by the Brown–Forsythe test. Group means of heteroscedastic data were then compared using Welch’s one-way analysis of variance (ANOVA) with a family-wise significance level of $\alpha=0.05$. Post-hoc pairwise comparisons against the control group were performed using Dunnett’s T3 test, and adjusted *P*-values are reported. For ternary compositional data, multivariate group differences were assessed by multivariate analysis of variance (MANOVA) on centered log-ratio (CLR)–transformed data, with statistical significance evaluated using the Hotelling–Lawley trace.

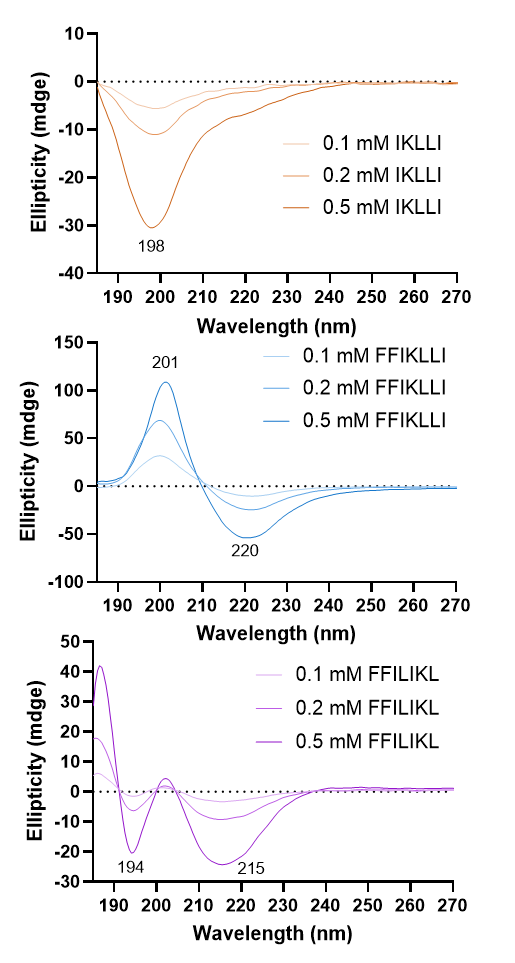

**Figure S1.** CD spectra of peptides **IKLLI**, **FFIKLLI**, and **FFILIKL** in water at different concentrations.

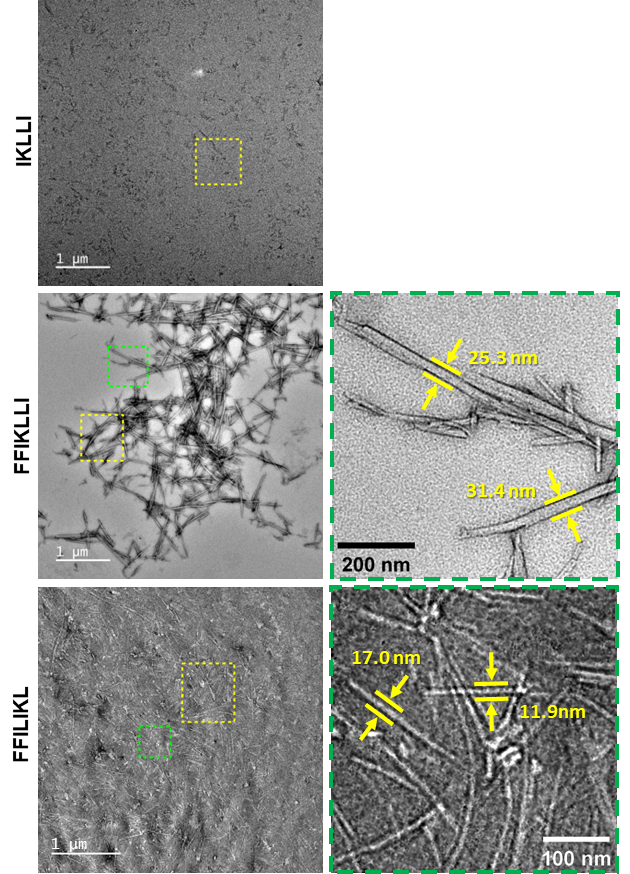

**Figure S2**. TEM images of **IKLLI**, **FFIKLLI**, and **FFILIKL** assemblies in water (10 mM). Yellow dashed boxes denote regions enlarged in Figure 1d. Green dashed boxes indicate regions enlarged to visualize fiber widths, which are labeled in the corresponding images.

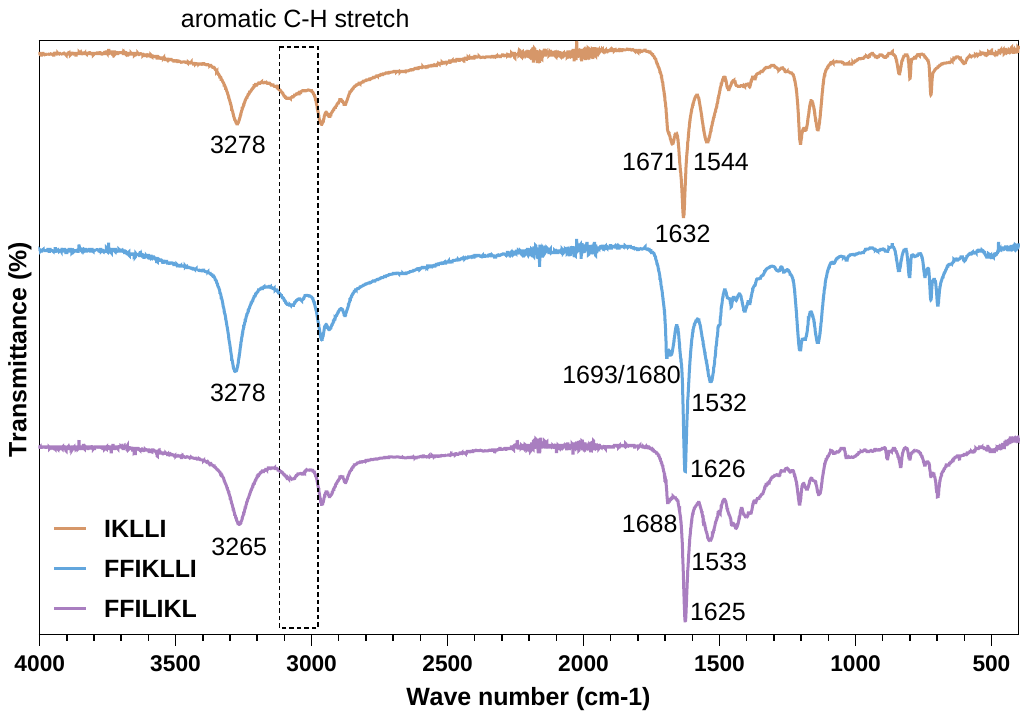

**Figure S3.** FTIR spectra of **IKLLI**, **FFIKLLI**, and **FFILIKL** assemblies.

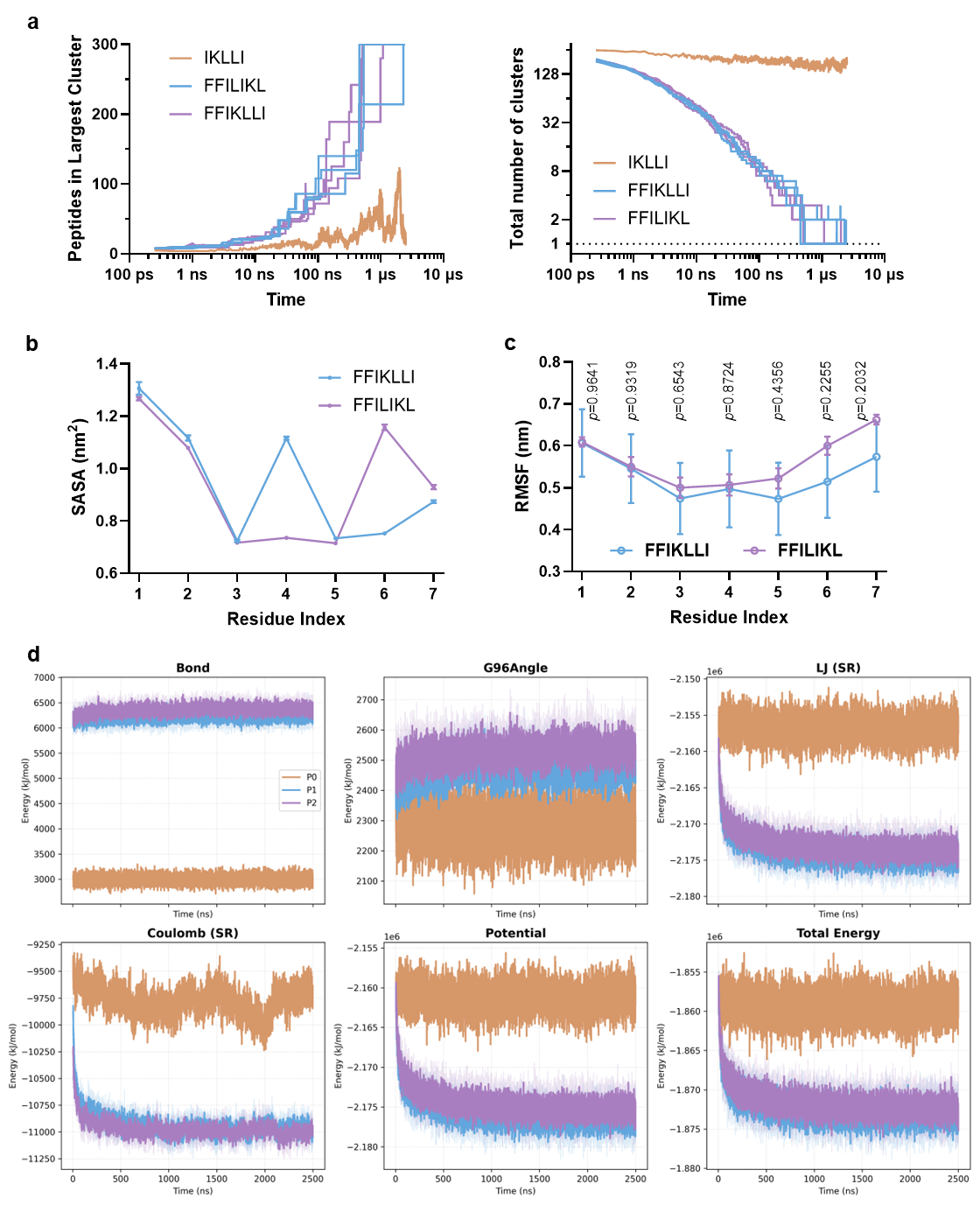

**Figure S4.** **Coarse-grained molecular dynamics (CG-MD) analysis of peptide self-assembly.** **(a)** Time evolution of peptide aggregation during CG-MD simulations. Maximum cluster size (left) and total cluster number (right) are shown as a function of simulation time for **IKLLI**, **FFIKLLI**, and **FFILIKL**. Whereas FF-containing peptides progressively consolidated into large aggregates, **IKLLI** remained distributed among multiple smaller clusters throughout the simulation. **(b)** Per-residue solvent-accessible surface area (SASA) of **FFIKLLI** and **FFILIKL** assemblies calculated from the final 20% of simulation trajectories after reaching equilibrium. Residue positions are shown from the N- to C-terminus. Lysine residues exhibited substantially higher solvent accessibility than neighboring hydrophobic residues, consistent with preferential surface exposure of charged ligand motifs. Data are presented as mean ± s.d. (n = 3 independent simulations) **(c)** Per-residue root mean square fluctuation (RMSF) of **FFIKLLI** and **FFILIKL** assemblies calculated from the final 20% of simulation trajectories. Residue positions are shown from the N- to C-terminus. No significant differences in residue-level mobility were detected between **FFIKLLI** and **FFILIKL** (two-tailed Welch’s t-test, p > 0.05). Data are presented as mean ± s.d. (n = 3 independent simulations). **(d)** Time evolution of bonded (Bond and G96Angle), non-bonded (LJ_SR and Coulomb_SR), total potential, and total energies during CG-MD simulations. Compared with **IKLLI**, FF-containing peptides exhibited lower non-bonded interaction energies and total potential energies, consistent with enhanced intermolecular packing during self-assembly. Despite their distinct sequence arrangements, **FFIKLLI** and **FFILIKL** displayed broadly similar energetic profiles, suggesting that incorporation of the FF assembly motif is a dominant contributor to bulk assembly energetics.

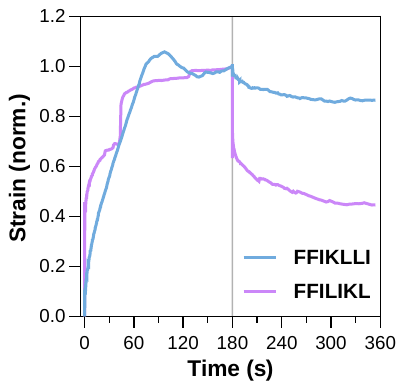

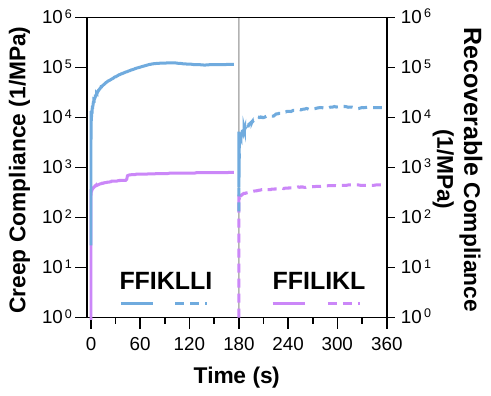

**a**

**b**

**Figure 5.** Strain **(a)** and compliance **(b)** profiles of **FFIKLLI** and **FFILIKL** assemblies obtained from creep-recovery measurements.

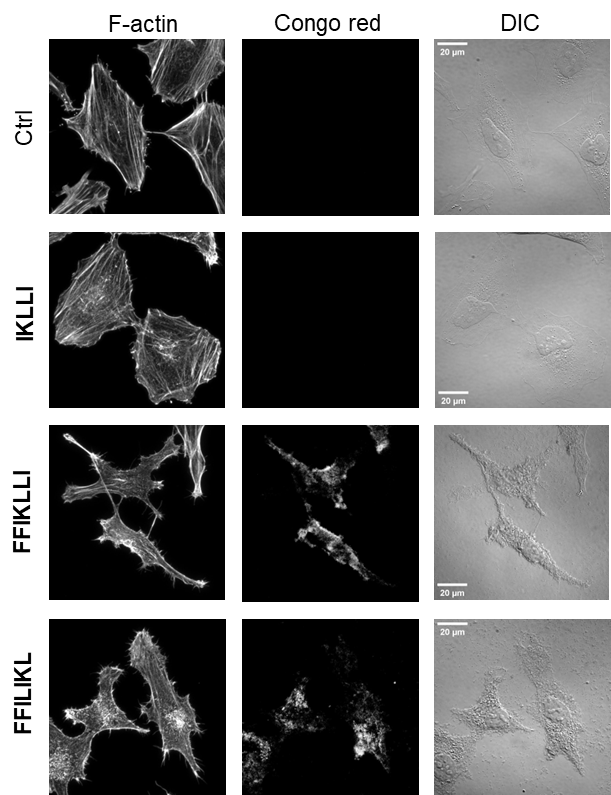

**Figure S6.** Individual channel images of F-actin and Congo red, presented as maximum intensity projections (MIPs), together with differential interference contrast (DIC) images, for the Ctrl, **IKLLI**, **FFIKLLI**, and **FFILIKL** groups, corresponding to Figure 2a.

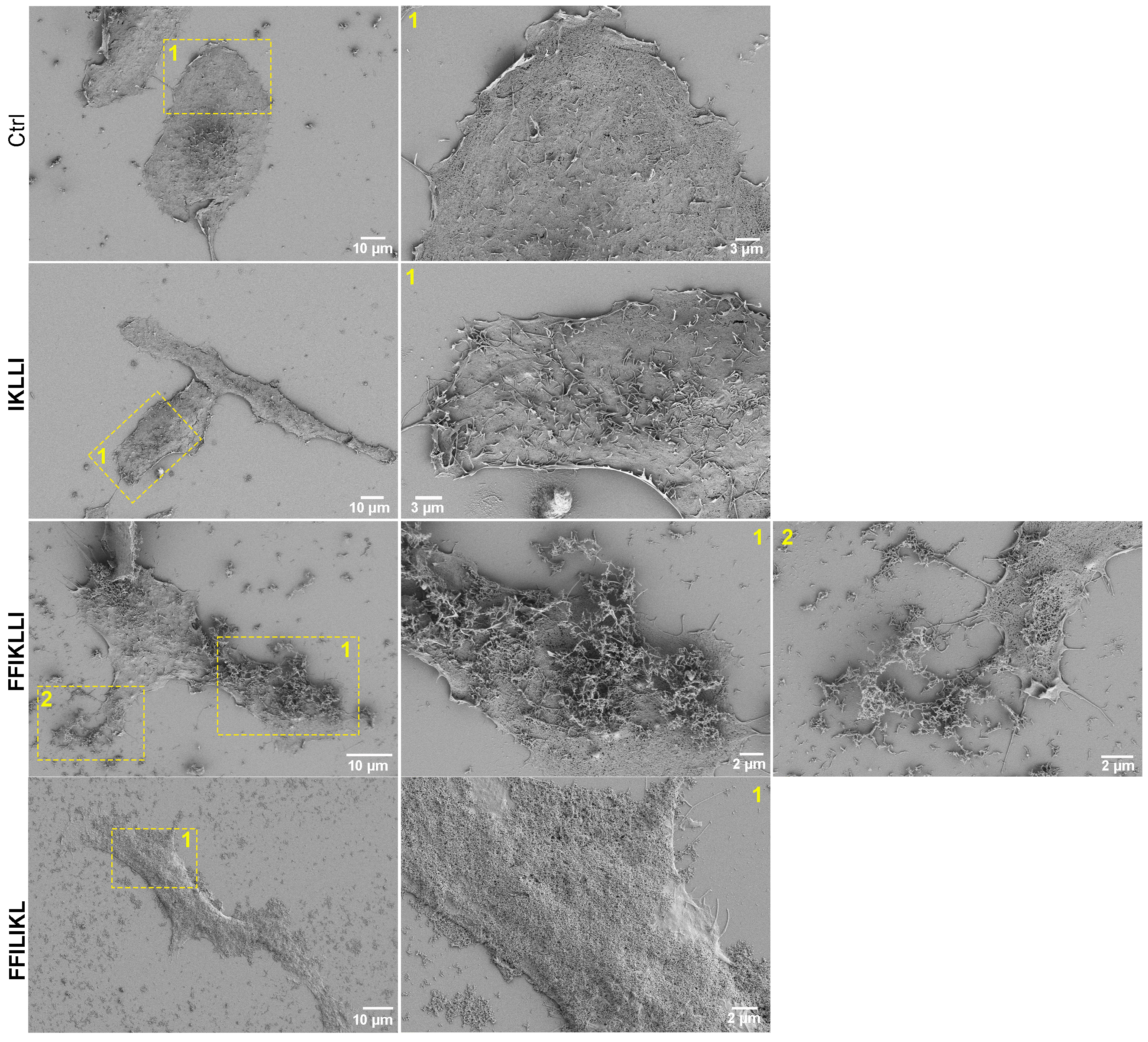

**Figure S7.** Scanning electron microscopy (SEM) images of untreated HeLa cells (Ctrl) and HeLa cells treated with **IKLLI**, **FFIKLLI**, or **FFILIKL** assemblies.

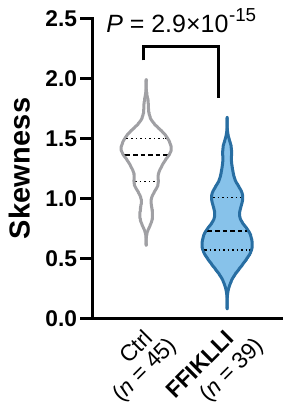

**Figure S8.** Skewness analysis of the distributions in figure 2i, and 2j quantifies the **FFIKLLI**-induced upward redistribution of activated integrin *β*1 along the Z-axis, reflected by altered distribution asymmetry between groups.

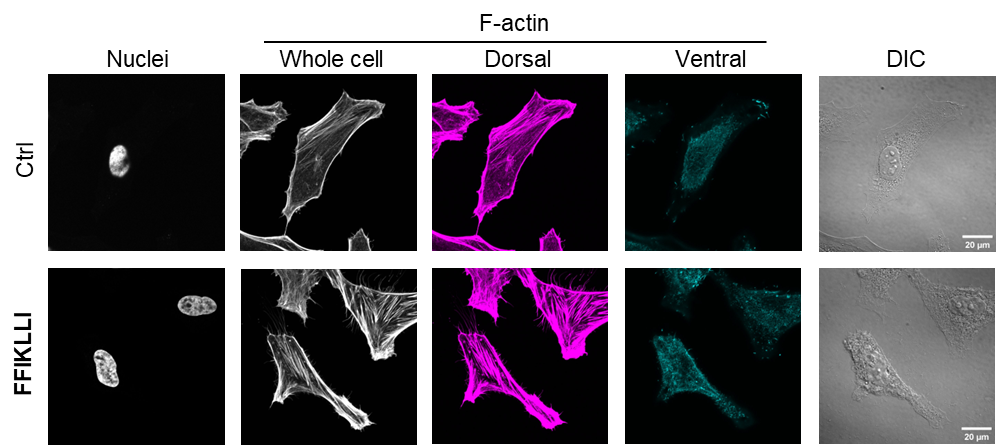

**Figure S9**. Individual channel images of nuclei, F-actin, and DIC for control and **FFIKLLI**-treated cells (100 μM, 24 h), corresponding to Figure 2g–l. F-actin is shown as maximum intensity projections (MIPs) generated from z-slices encompassing the whole cell (gray), dorsal compartment (magenta), or ventral compartment (cyan). Different lookup tables (LUTs) and display ranges were applied to compartment-specific and whole-cell projections to facilitate visualization of lower-intensity dorsal F-actin structures.

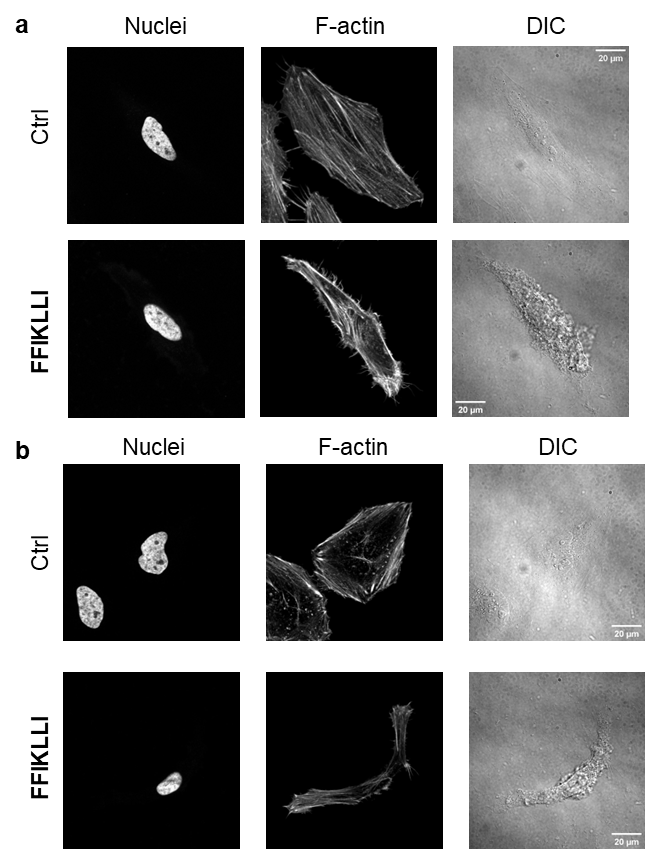

**Figure S10.** Individual channel images of nuclei, F-actin, and DIC for control and **FFIKLLI**-treated cells (100 μM, 24 h), corresponding to Figure 2m. In pMLC-stained cells (a), F-actin is shown as maximum intensity projections (MIPs) of five consecutive z-slices acquired at 0.5 μm intervals and centered on the ventral stress fiber–rich plane. In paxillin-stained cells (b), F-actin is shown as a representative single z-slice corresponding to the focal adhesion–rich ventral plane.

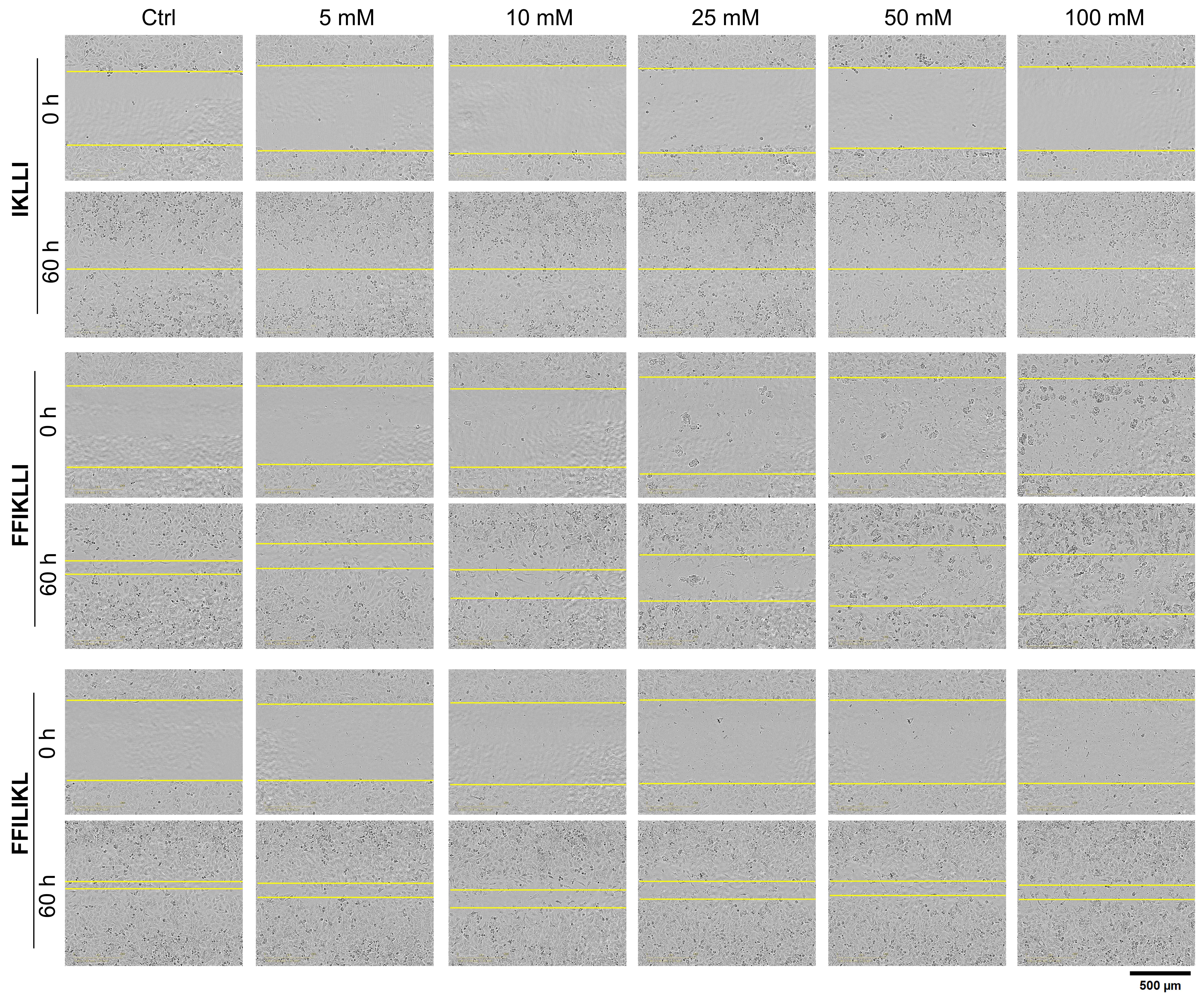

**Figure S11**. Representative bright-field images from wound-healing assays of HeLa cells treated with **IKLLI**, **FFIKLLI**, or **FFILIKL** at five different concentrations, together with untreated controls, corresponding to Figure 3a.

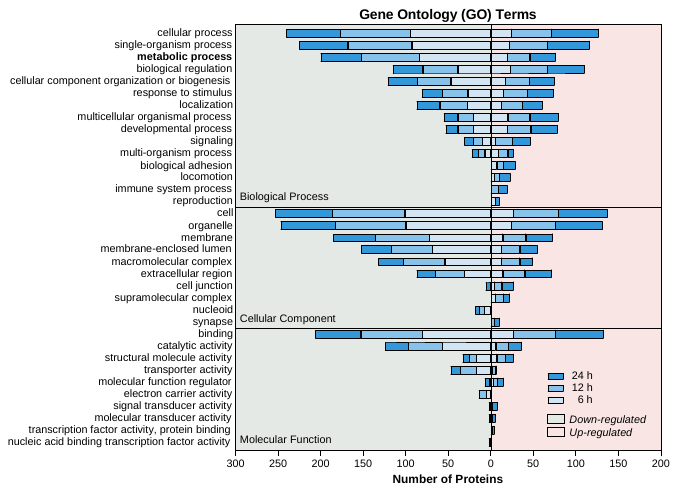

**Figure S12**. Gene Ontology (GO) annotation of proteins differentially expressed following treatment with 50 μM **FFIKLLI** for 6, 12, and 24 h. A total of 695 differentially expressed proteins were classified into the three major GO categories: biological process, cellular component, and molecular function, together with their enriched subcategories.

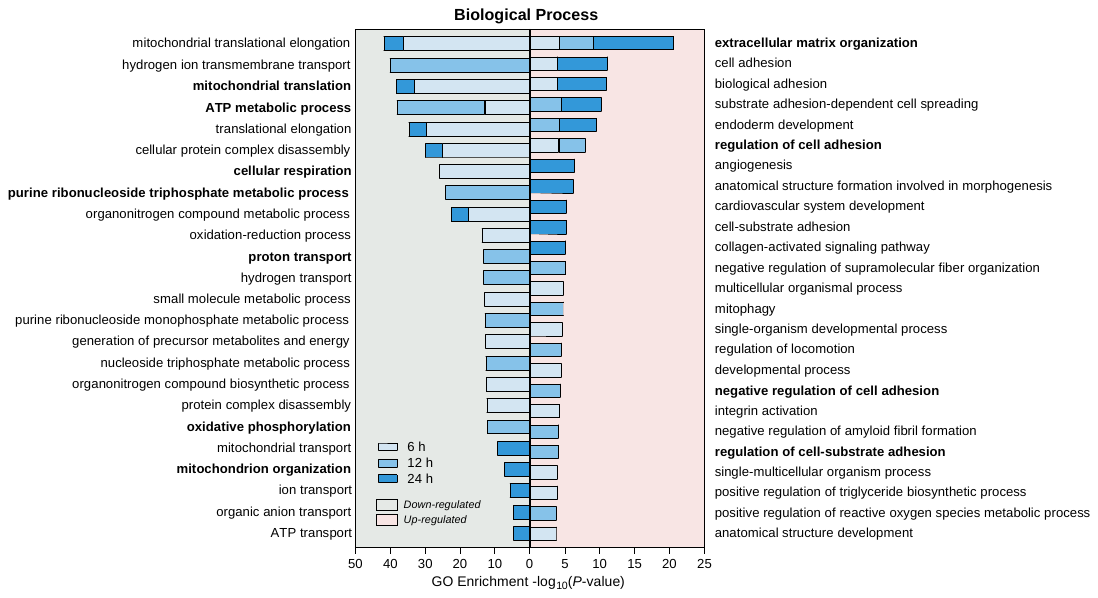

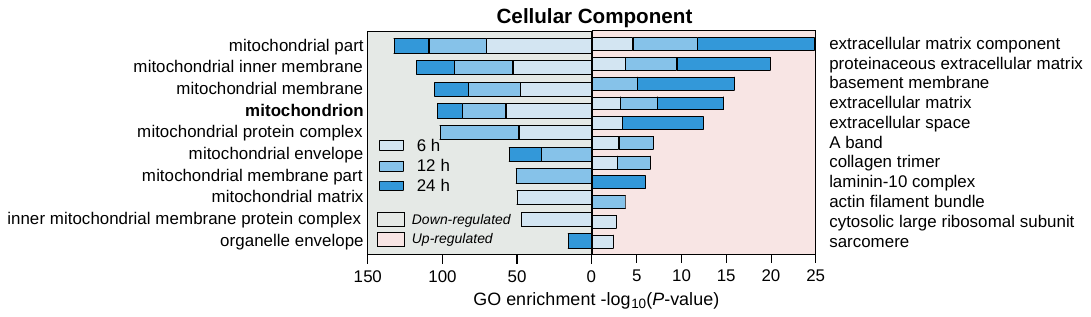

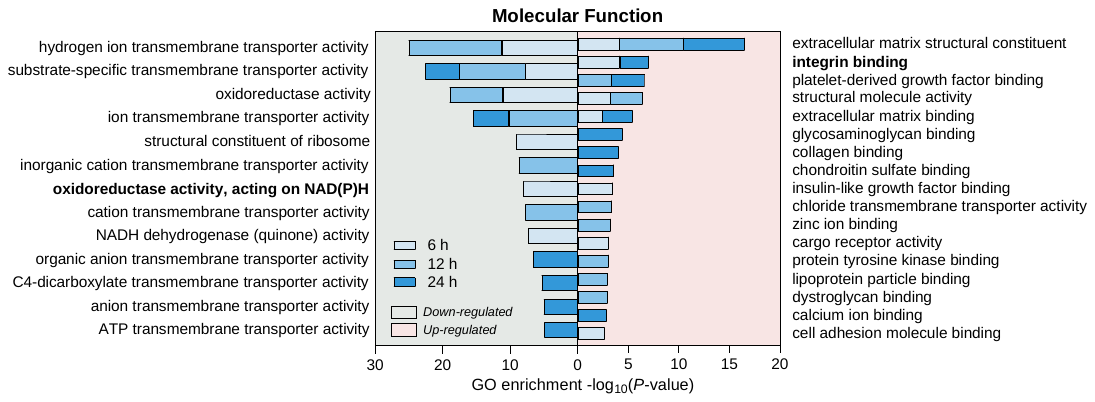

**Figure S13**. GO enrichment analysis of biological process, cellular component, and molecular function. The enrichment of significantly differentially expressed proteins in each GO subcategory relative to all identified proteins was assessed by two-tailed Fisher's exact test and presented as −*log*_10_(*P*-value) for down-regulated (left panel) and up-regulated (right panel) proteins.

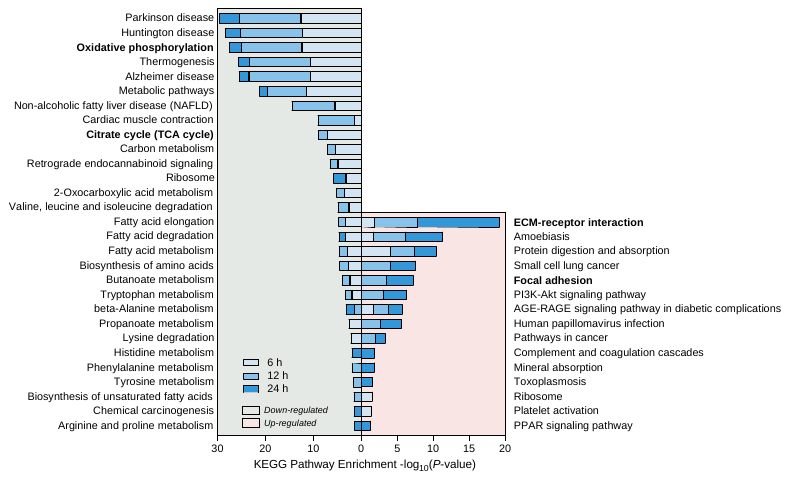

**Figure S14.** KEGG pathway enrichment analysis of significantly differentially expressed proteins relative to all identified proteins was assessed by two-tailed Fisher's exact test and presented as −*log*_10_(*P*-value) for down-regulated (left panel) and up-regulated (right panel) proteins. Oxidative phosphorylation as well as TCA cycle in mitochondria were significantly down-regulated.

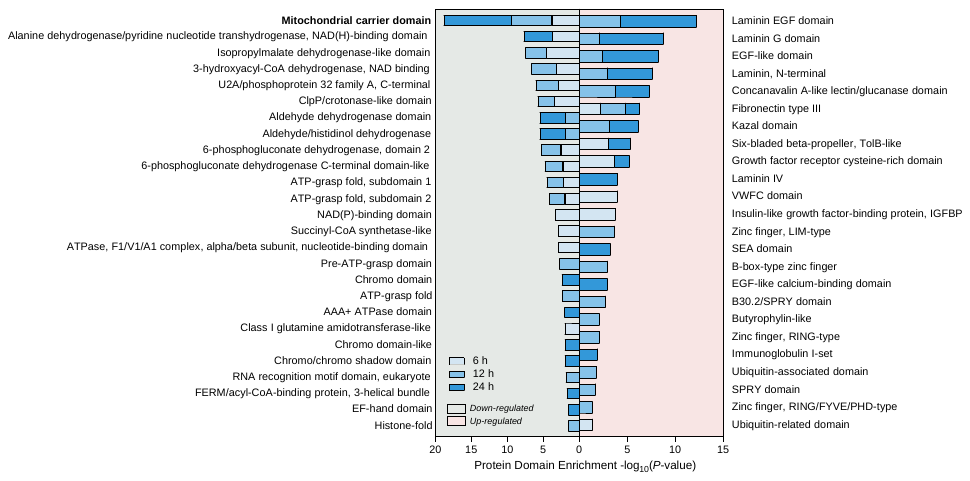

**Figure S15.** Protein domain enrichment analysis of significantly differentially expressed proteins relative to all identified proteins was assessed by two-tailed Fisher's exact test and presented as −*log*_10_(*P*-value) for down-regulated (left panel) and up-regulated (right panel) proteins. Mitochondrial carrier domain was significantly down-regulated.

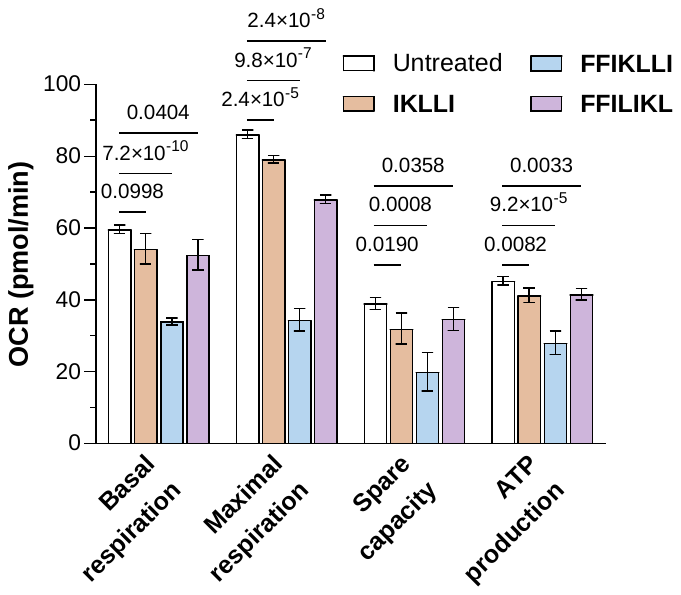

**Figure S16**. Mitochondrial respiration analysis in HeLa cells **following peptide treatment (100 μM, 12 h) using the** mitochondrial stress test, corresponding to Figure 3j. Values are mean ± s.d. (*n* = 5). Group differences were assessed by Brown–Forsythe and Welch ANOVA followed by Dunnett’s T3 post-hoc comparisons versus the untreated group. Adjusted *P*-values are shown.

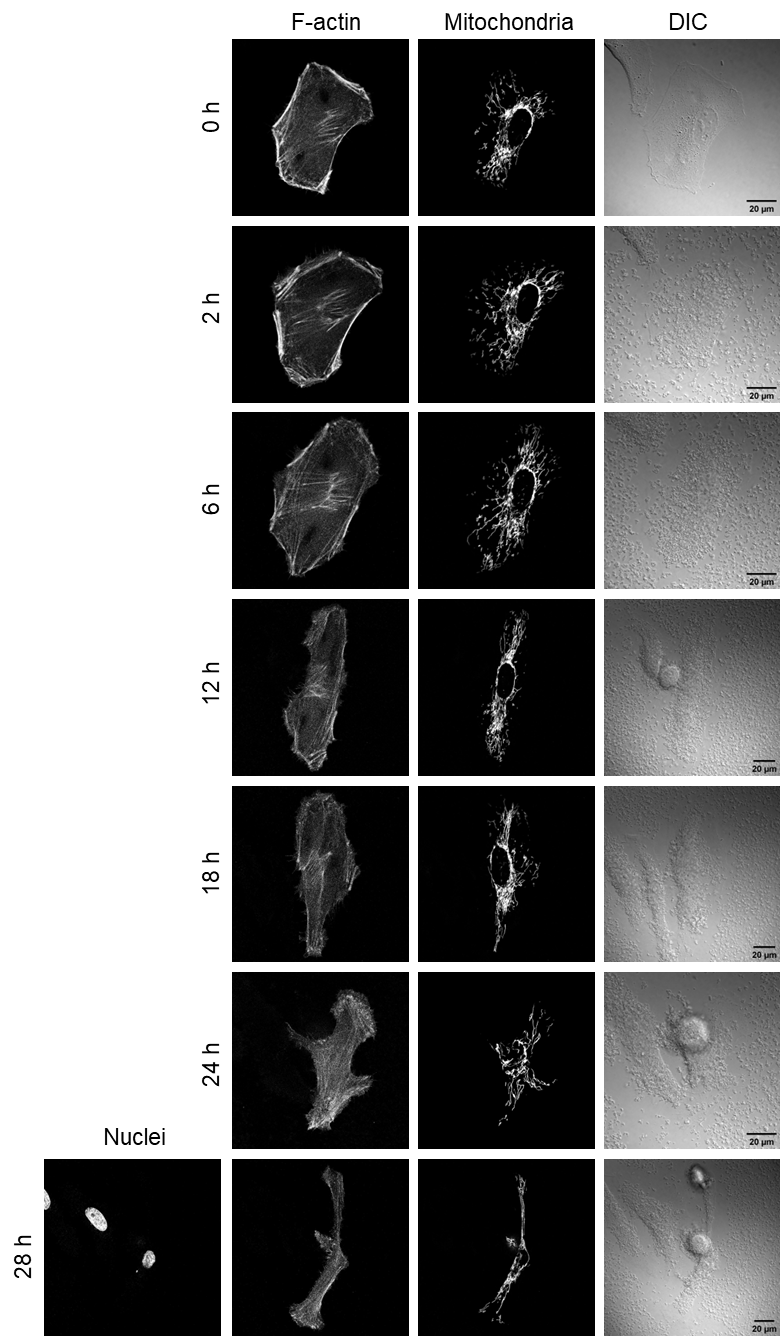

**Figure S17**. Individual channel images of F-actin, mitochondria, and DIC in live HeLa cells following treatment with 100 μM **FFIKLLI**, corresponding to Figure 4a.

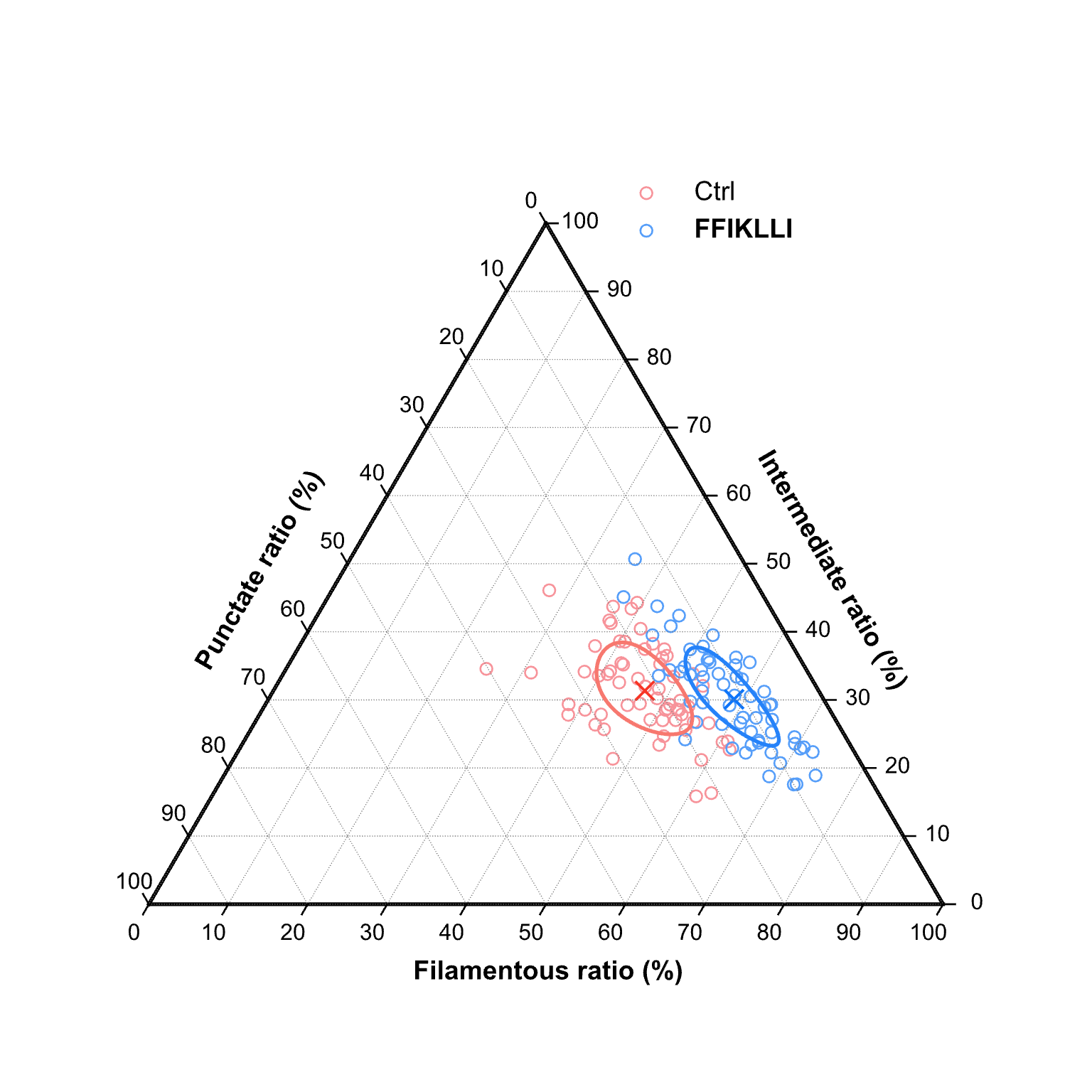

**Figure S18.** Ternary plot of mitochondrial morphology composition, expressed as the area fractions of punctate, intermediate, and filamentous mitochondria. Each hollow circle represents an individual cell, and crosses (×) indicate group means. Solid curves represent 1σ contours derived in centered log-ratio (CLR) space and projected onto the simplex. Statistical significance was determined using the Hotelling–Lawley trace from multivariate analysis of variance (MANOVA) of CLR-transformed compositional data.

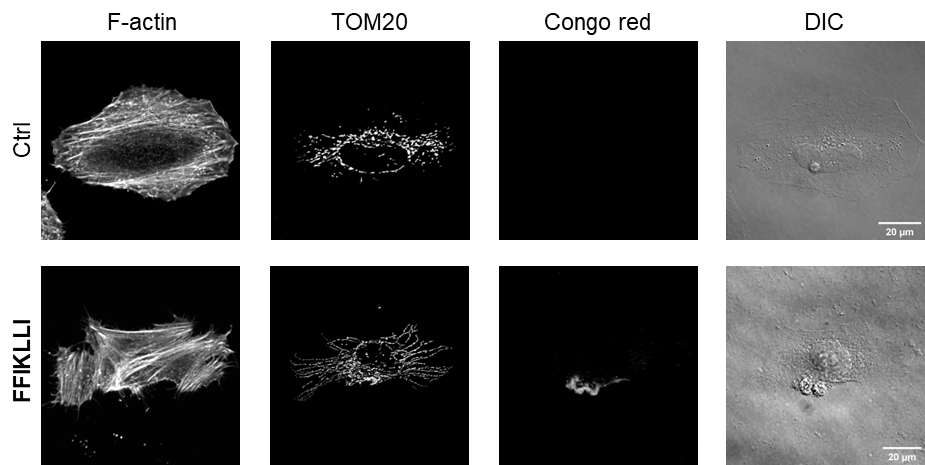

**Figure S19.** Individual channel images of F-actin, TOM20, Congo red, and DIC for control and **FFIKLLI**-treated cells (100 μM, 24 h), corresponding to Figure 4b–d.

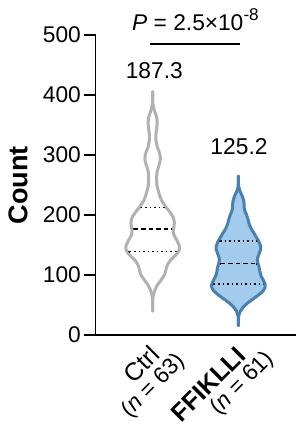

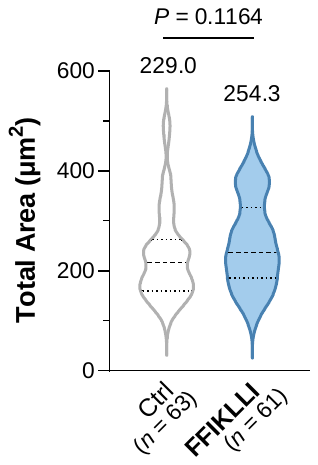

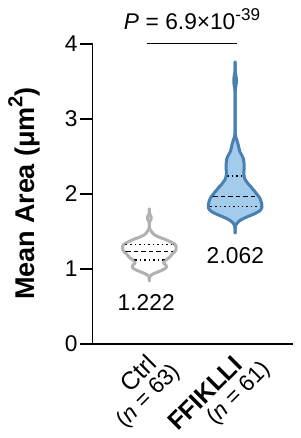

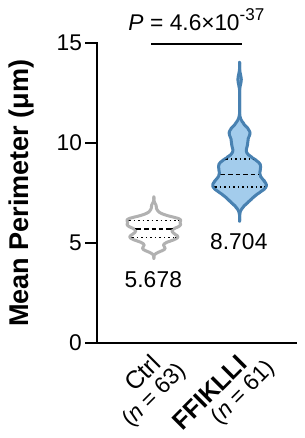

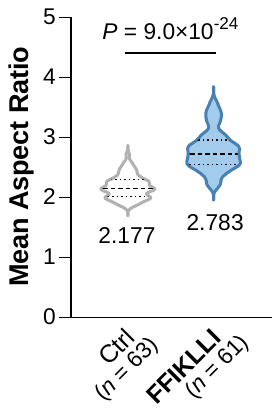

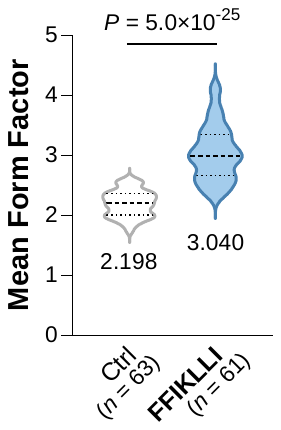

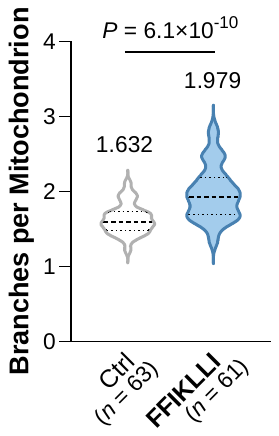

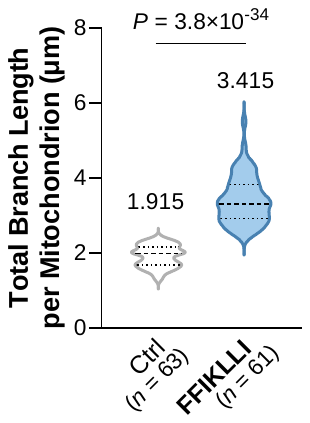

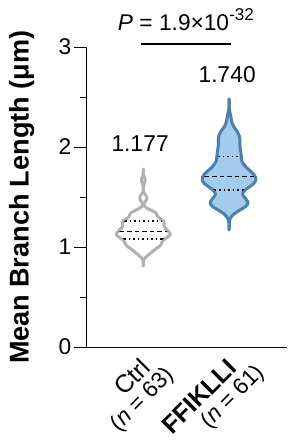

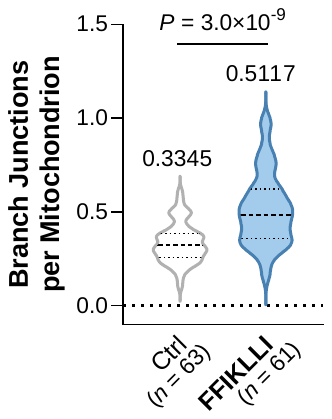

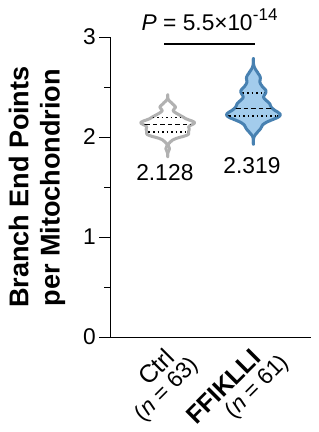

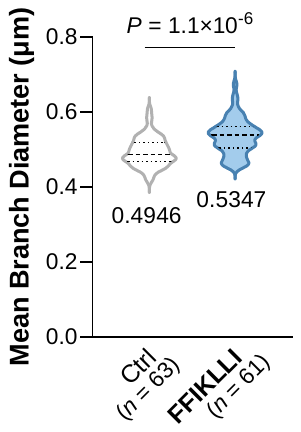

**Figure S20.** Mitochondrial morphometric analysis of control and **FFIKLLI**-treated cells (100 μM, 24 h), corresponding to Figure 4b–d. Mean values are annotated on each violin plot. Statistical significance was determined using a two-tailed Student's *t*-test.

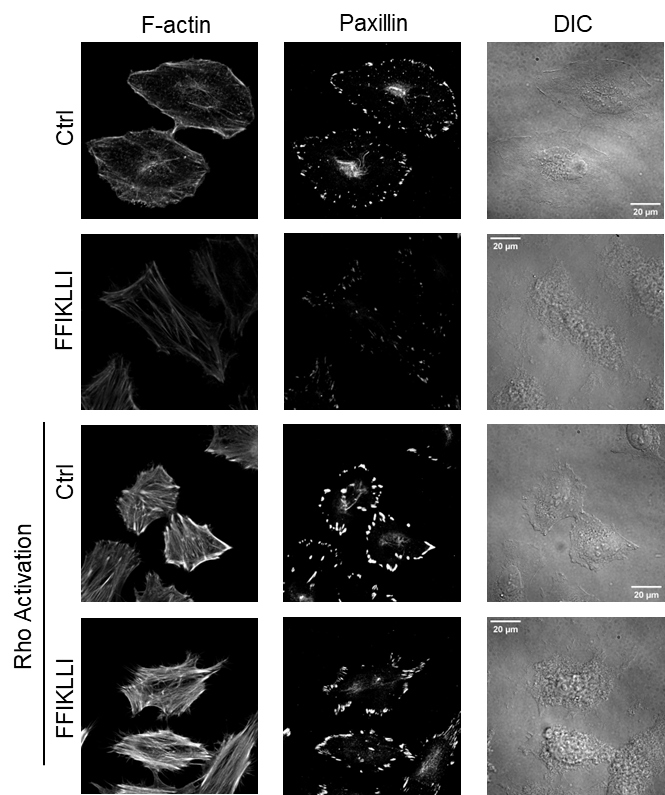

**Figure S21.** Individual channel images of F-actin, paxillin, and DIC from control and **FFIKLLI**-treated cells (100 μM, 24 h), with subsequent Rho Activator II treatment (4 h) where indicated, corresponding to Figure 4g.

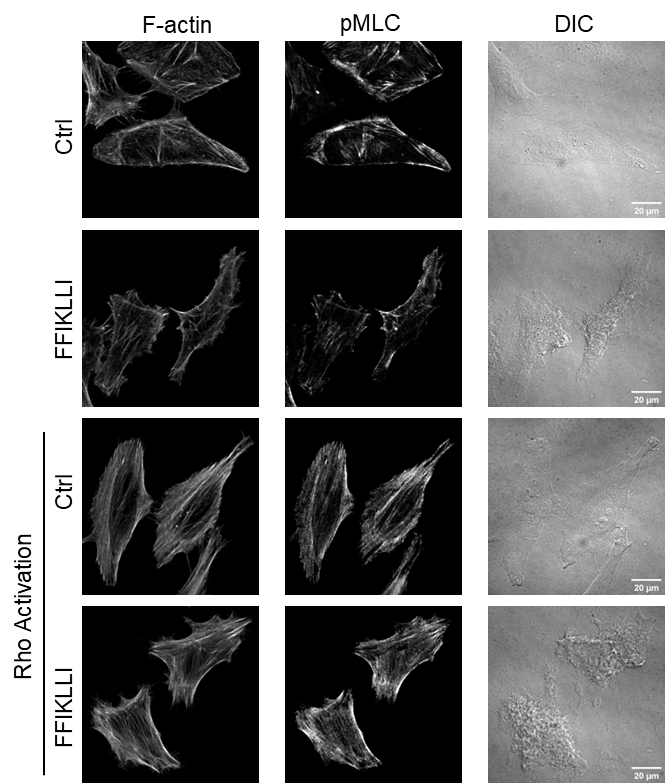

**Figure S22.** Individual channel images of F-actin, pMLC, and DIC for control and **FFIKLLI**-treated cells (100 μM, 24 h), corresponding to Figure 4g. F-actin and pMLC signals are shown as maximum intensity projections (MIPs) generated from five consecutive z-slices acquired at 0.5 μm intervals and centered on the ventral stress fiber–rich plane.

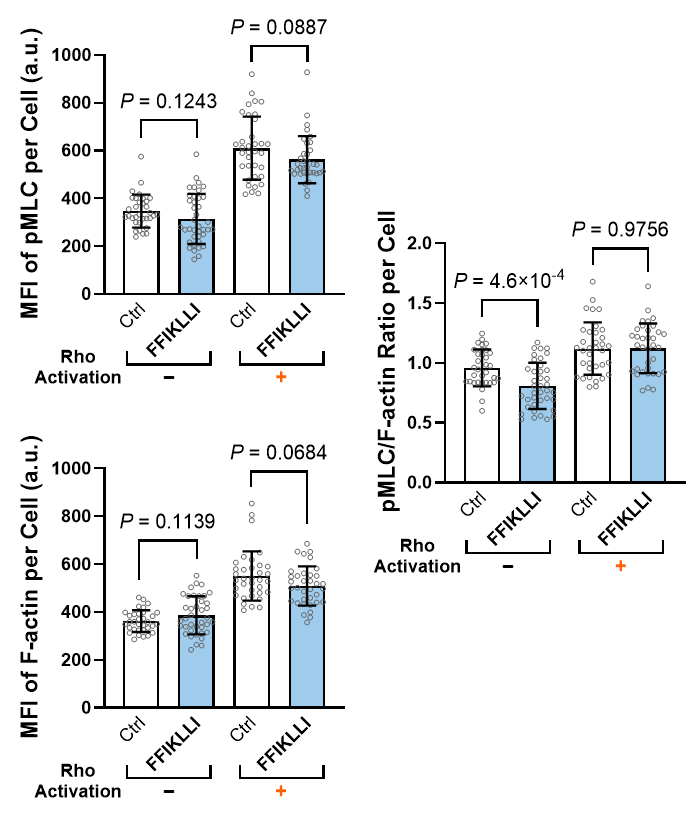

**Figure S23**. Quantification of pMLC and F-actin mean fluorescence intensities (MFIs), together with the pMLC-to-F-actin MFI ratio on a per-cell basis, corresponding to (4g). Data are presented as mean ± s.d. Statistical significance was determined using a two-tailed Welch's t-test. n = 33, 39, 35, and 35 cells.

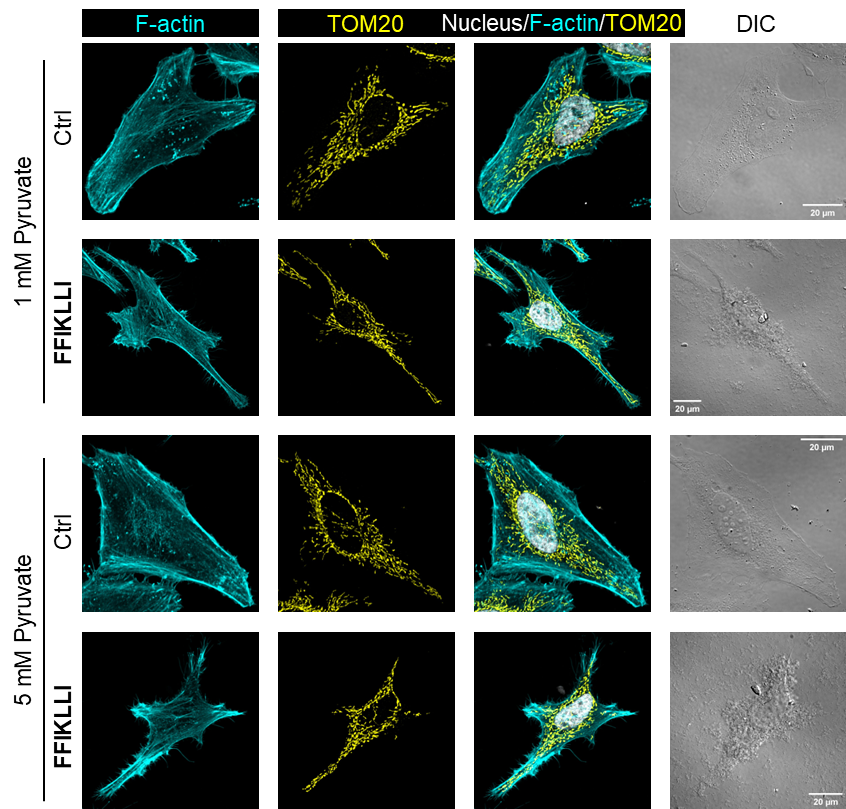

**Figure S24.** Confocal images of F-actin, TOM20, and DIC from control and **FFIKLLI**-treated cells (100 μM, 12 h), under normal (1 mM) or high (5 mM) pyruvate concentration where indicated, corresponding to Figure 4l–m.

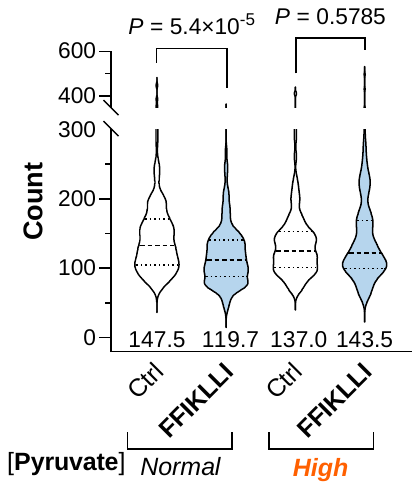

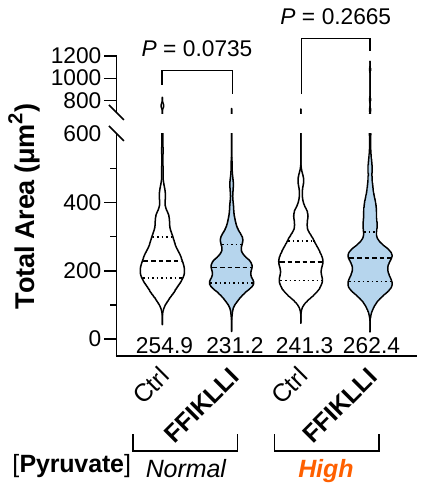

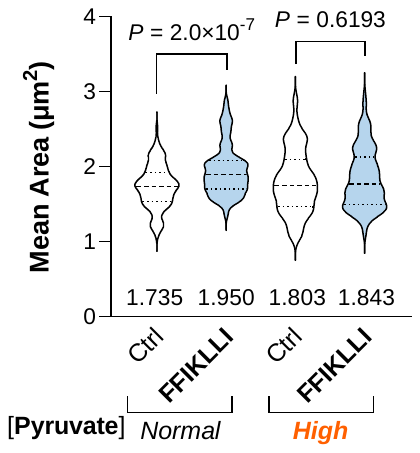

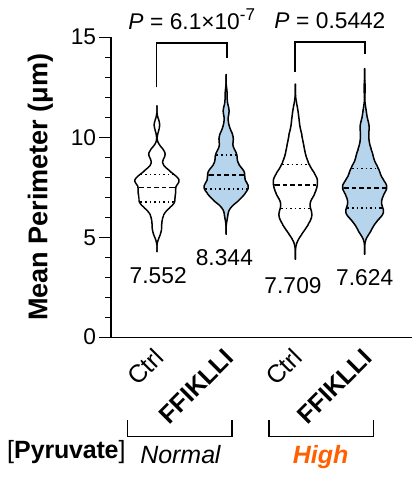

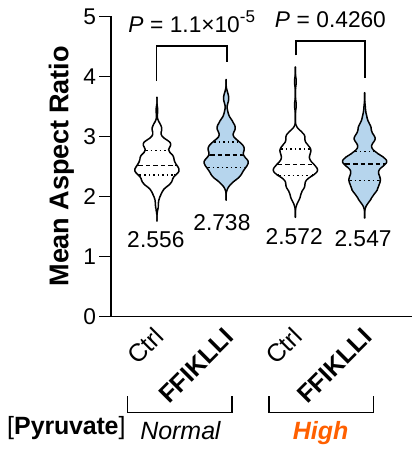

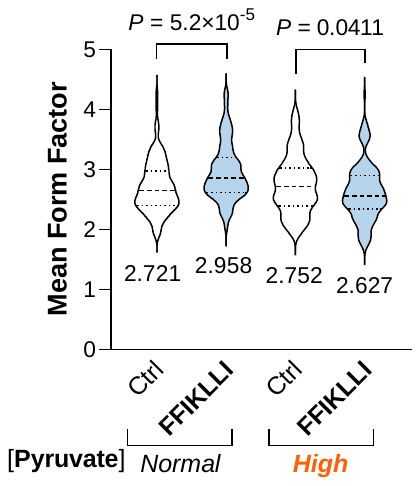

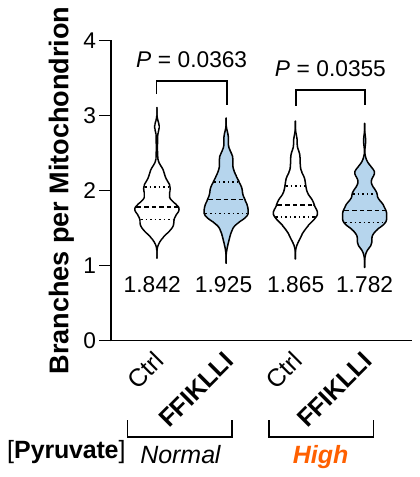

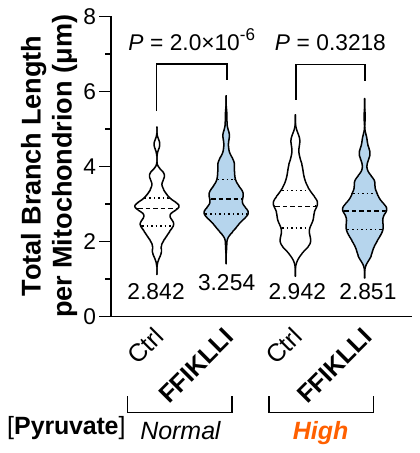

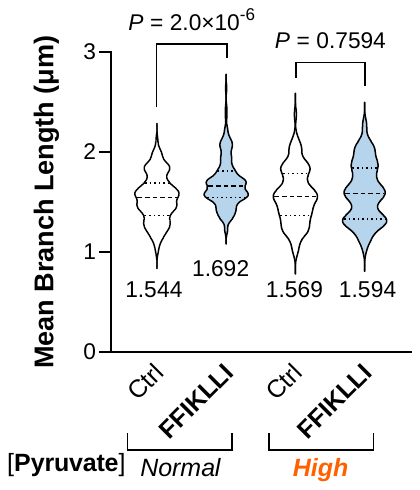

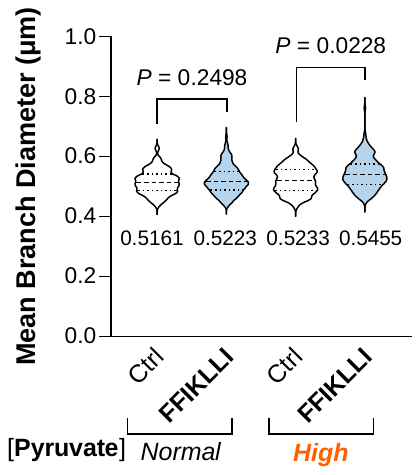

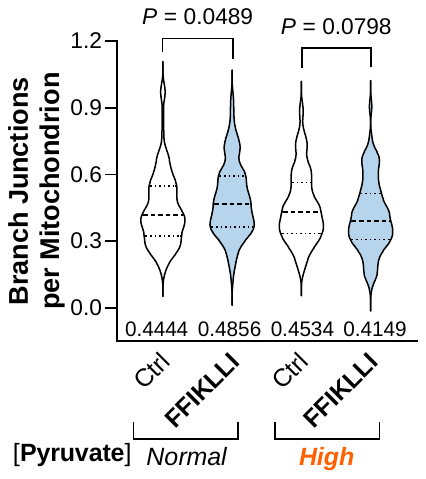

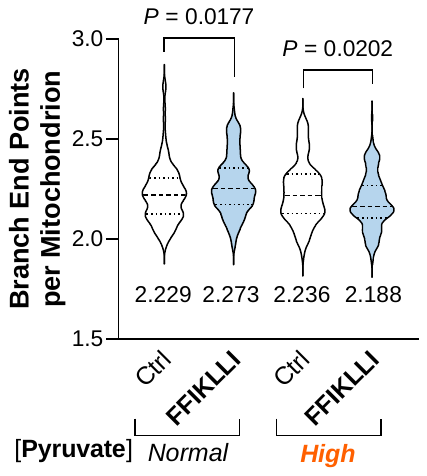

**Figure S25.** Mitochondrial morphometric analysis of control and **FFIKLLI**-treated cells (100 μM, 12 h) under normal (1 mM) or high (5 mM) pyruvate concentration, corresponding to Figure S24. Mean values are annotated on each violin plot. Statistical significance was determined using a two-tailed Student's *t*-test.

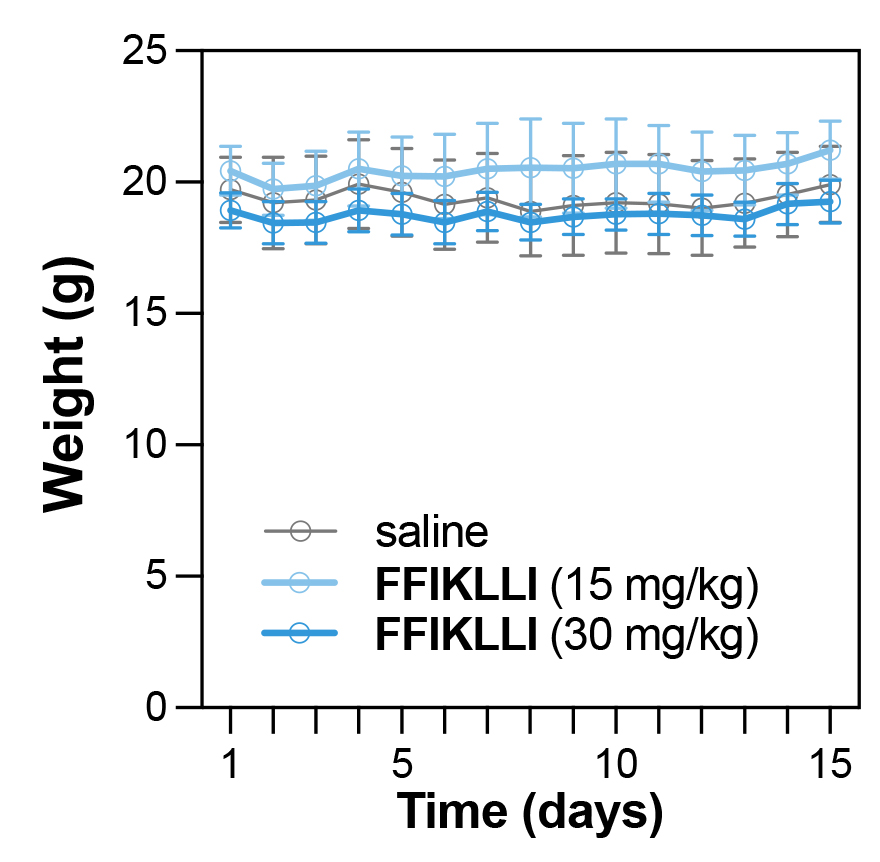

**Figure S26.** Body weight changes of mice throughout the 15-day HeLa xenograft study. Data are presented as mean ± s.d. (n = 5 mice per group).

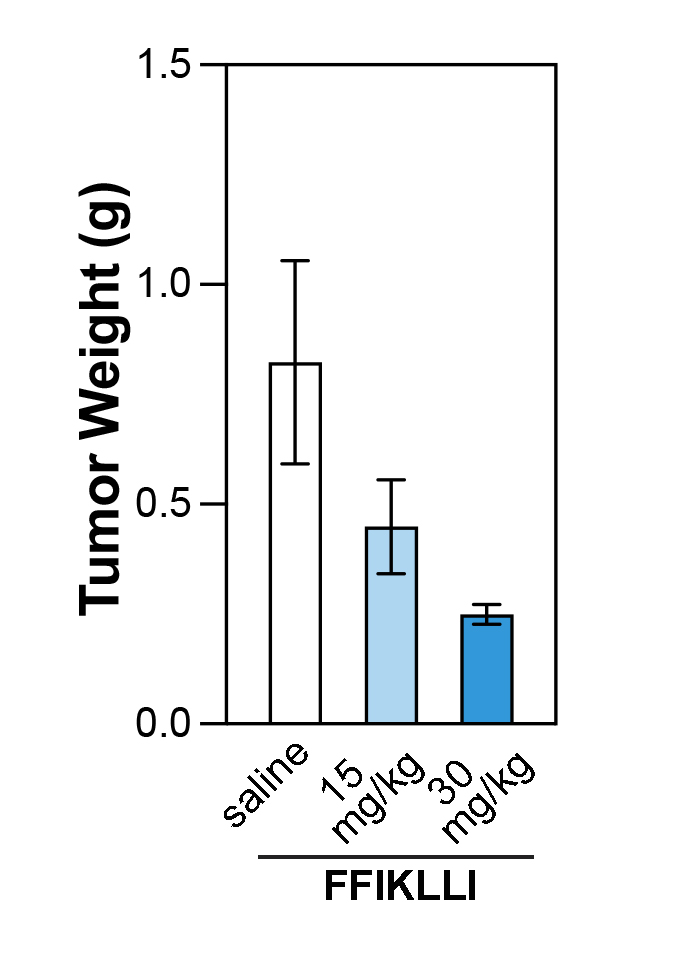

**Figure S27.** Excised tumor weights at the end of the 15-day HeLa xenograft study. Data are presented as mean ± s.d. (n = 5 mice per group).

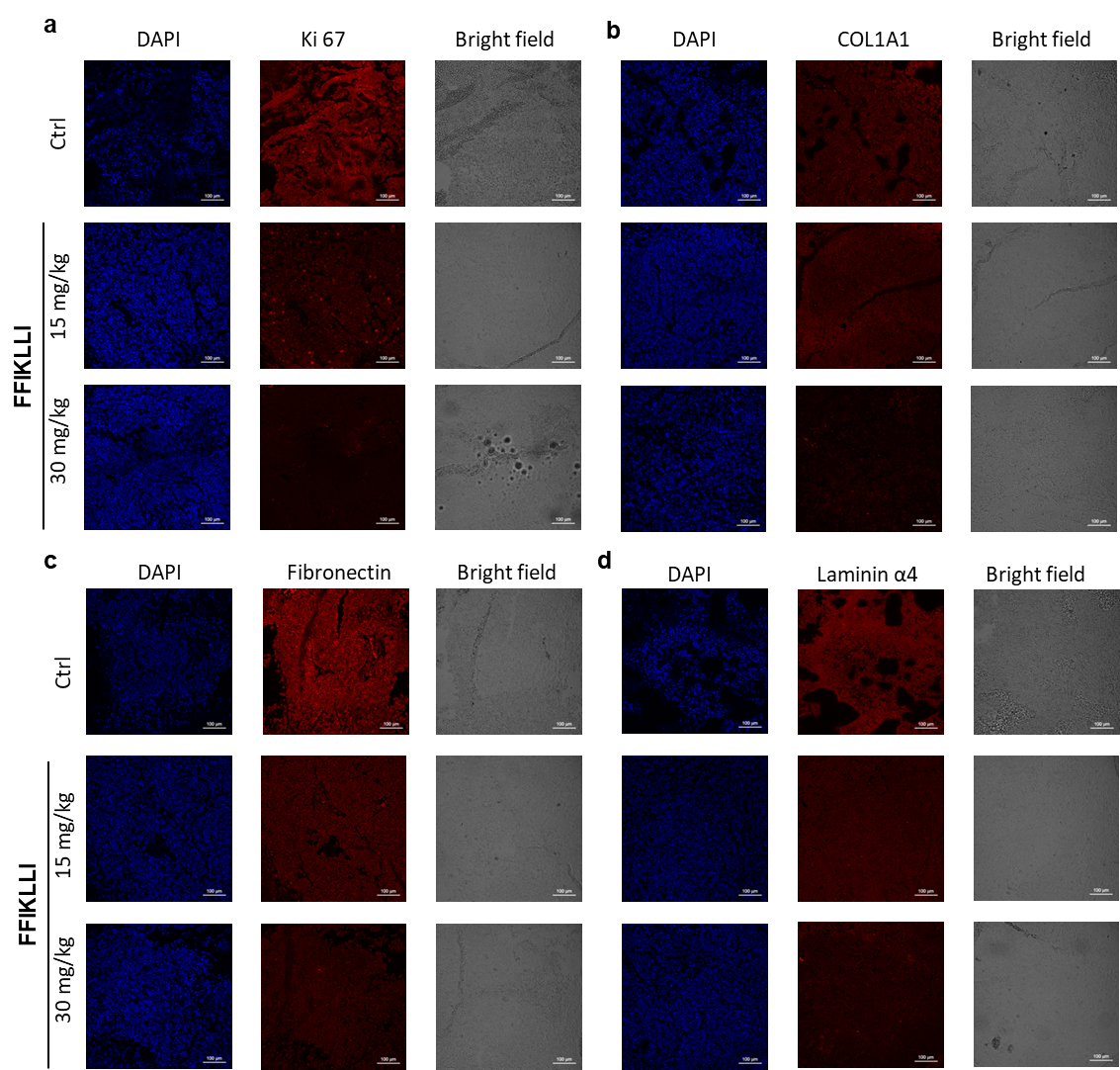

**Figure S28.** Individual channel images corresponding to the merged immunofluorescence images presented in Figure 5e,f. For each treatment group (vehicle control and **FFIKLLI** at 15 and 30 mg kg⁻¹) and immunostaining marker Ki67 (a), COL1A1 (b), laminin α4 (c), and fibronectin (d), the DAPI, immunofluorescence, and brightfield channels are displayed separately.

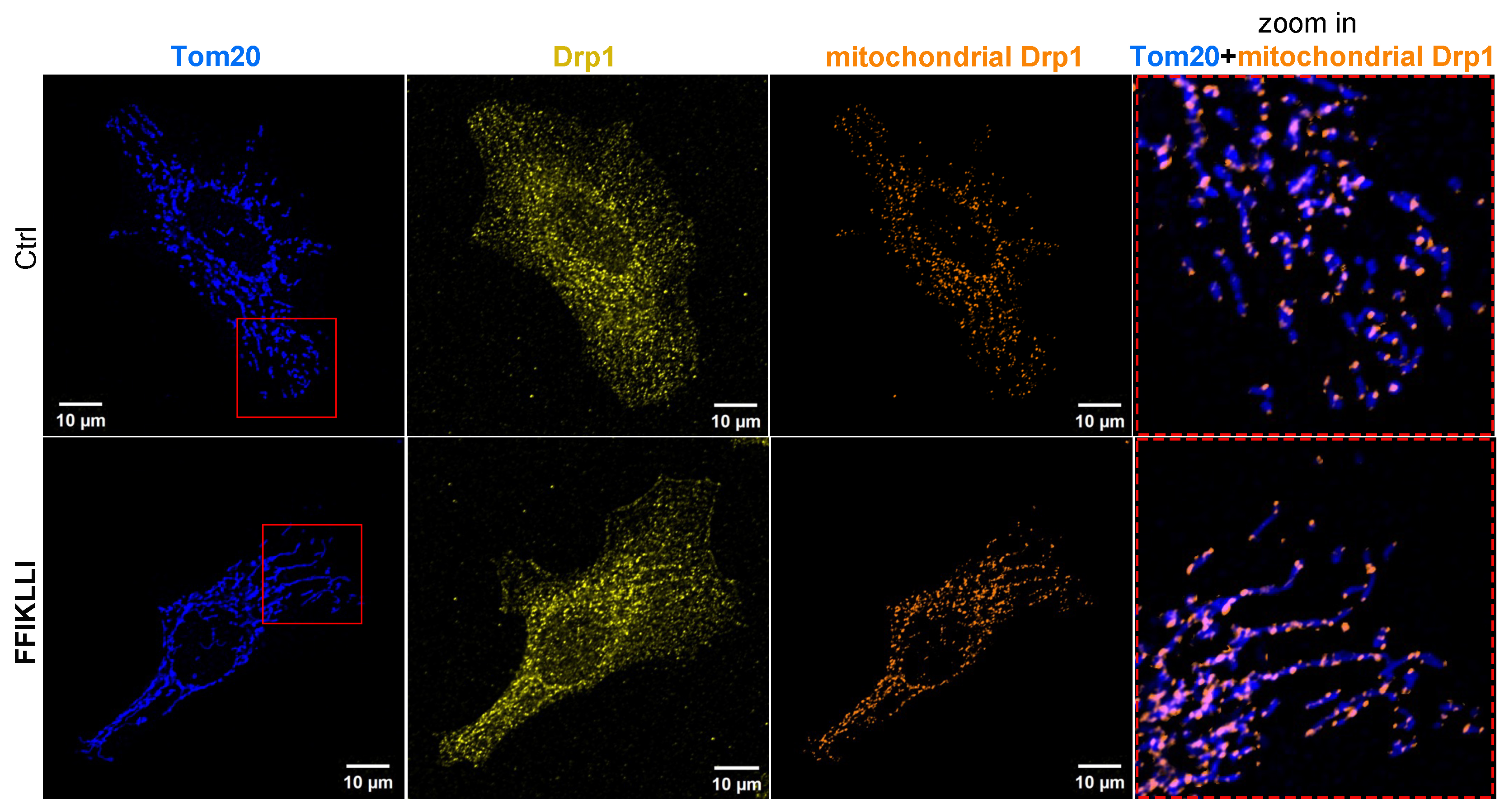

**Figure S29.** Representative immunofluorescence images of mitochondrial DRP1 localization in HeLa cells with or without **FFIKLLI** treatment. Mitochondria were labeled with TOM20 (blue), DRP1 is shown in yellow, and TOM20-colocalized DRP1 (mitochondrial DRP1) is displayed in orange. Insets show enlarged views of the merged TOM20 and mitoDRP1 signals.

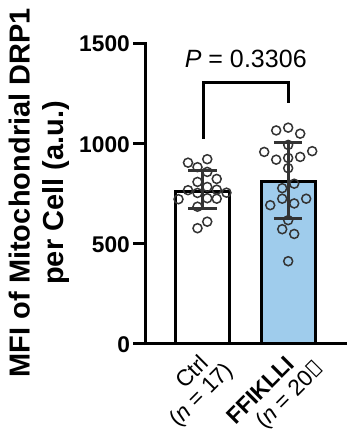

**Figure S30**. Quantification of mitochondria-associated DRP1 (mitochondrial DRP1) mean fluorescence intensity (MFI) per cell in HeLa cells treated with or without **FFIKLLI**. Two-tailed Welch's t-test was used to compare groups.

### Attachment 1. LC-MS spectra of peptide FFILIKL.

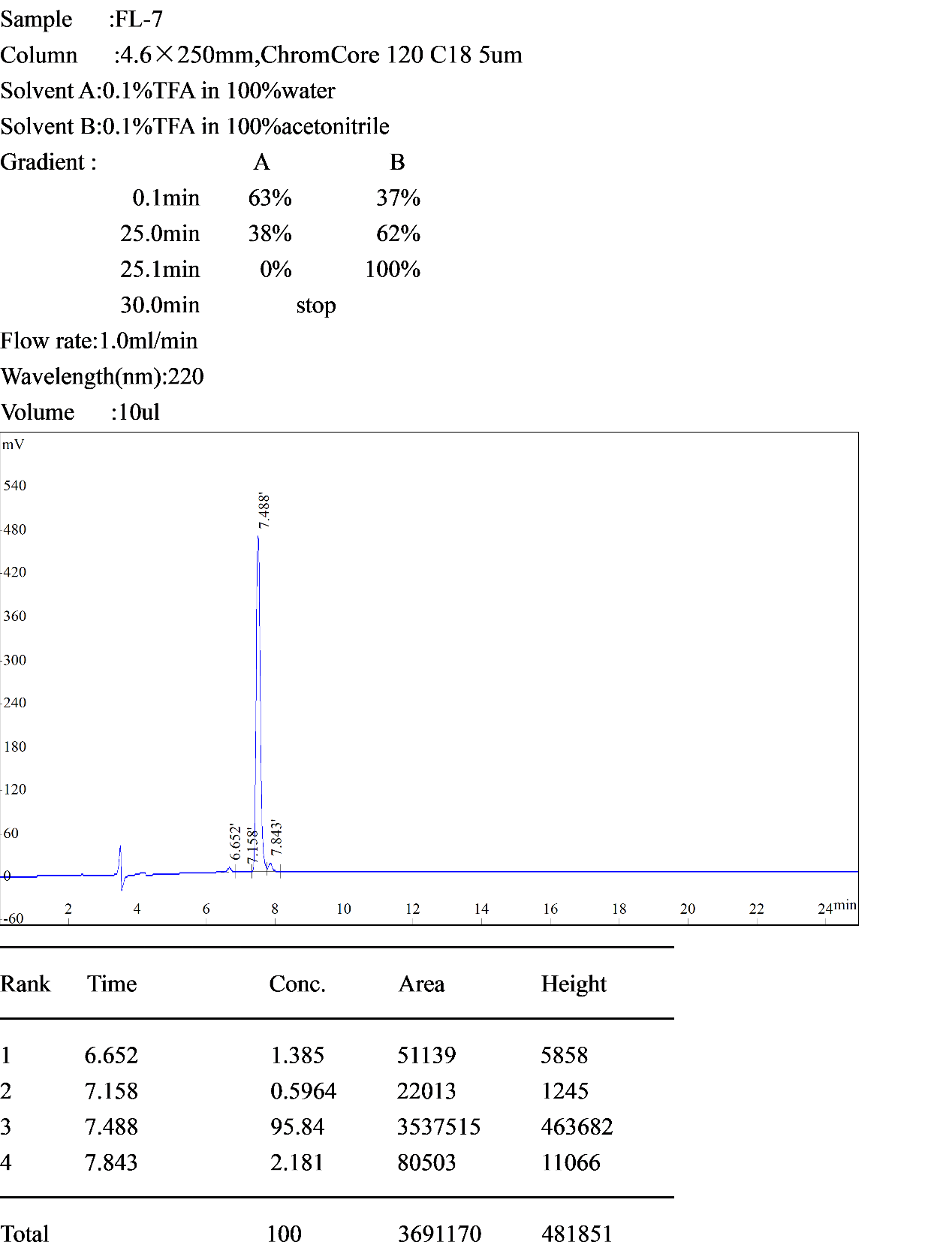

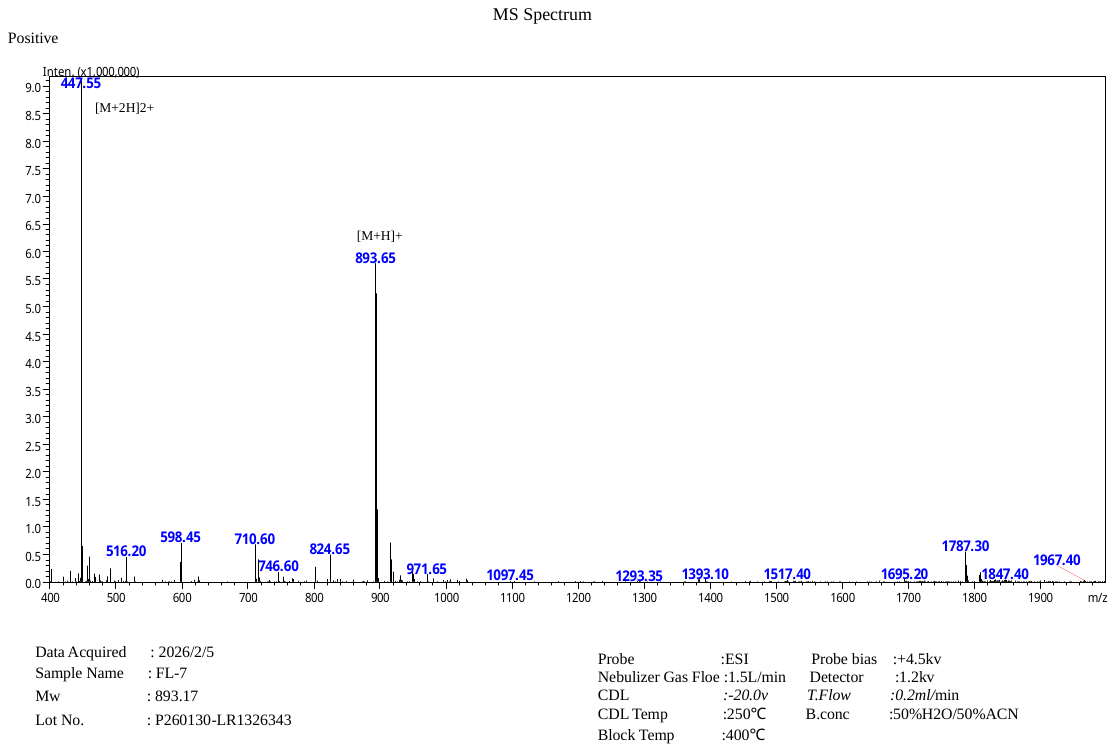

### Attachment 2. ^1^H NMR spectrum of peptide FFILIKL.

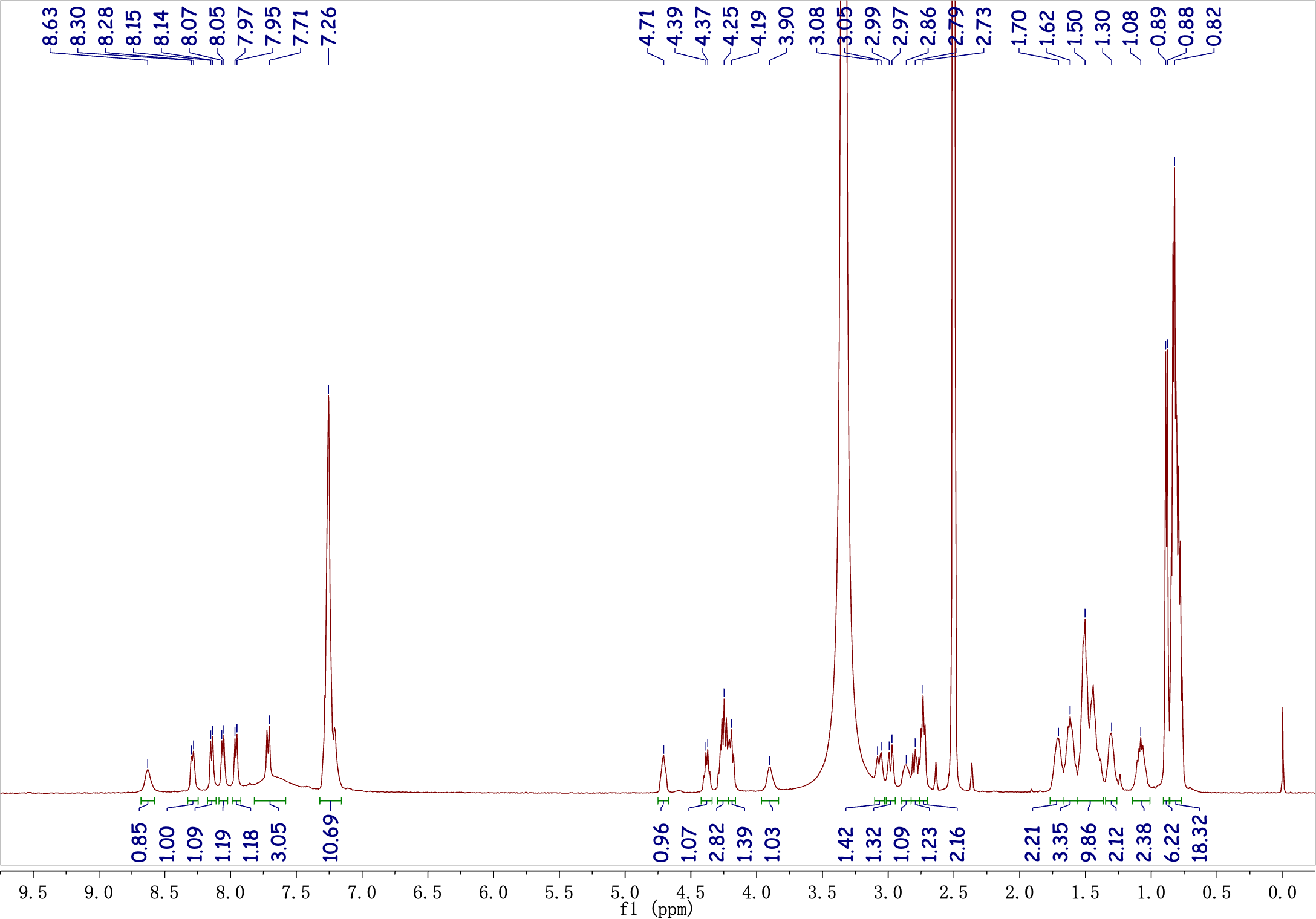

### Attachment 3. ^13^C NMR spectrum of peptide FFILIKL.

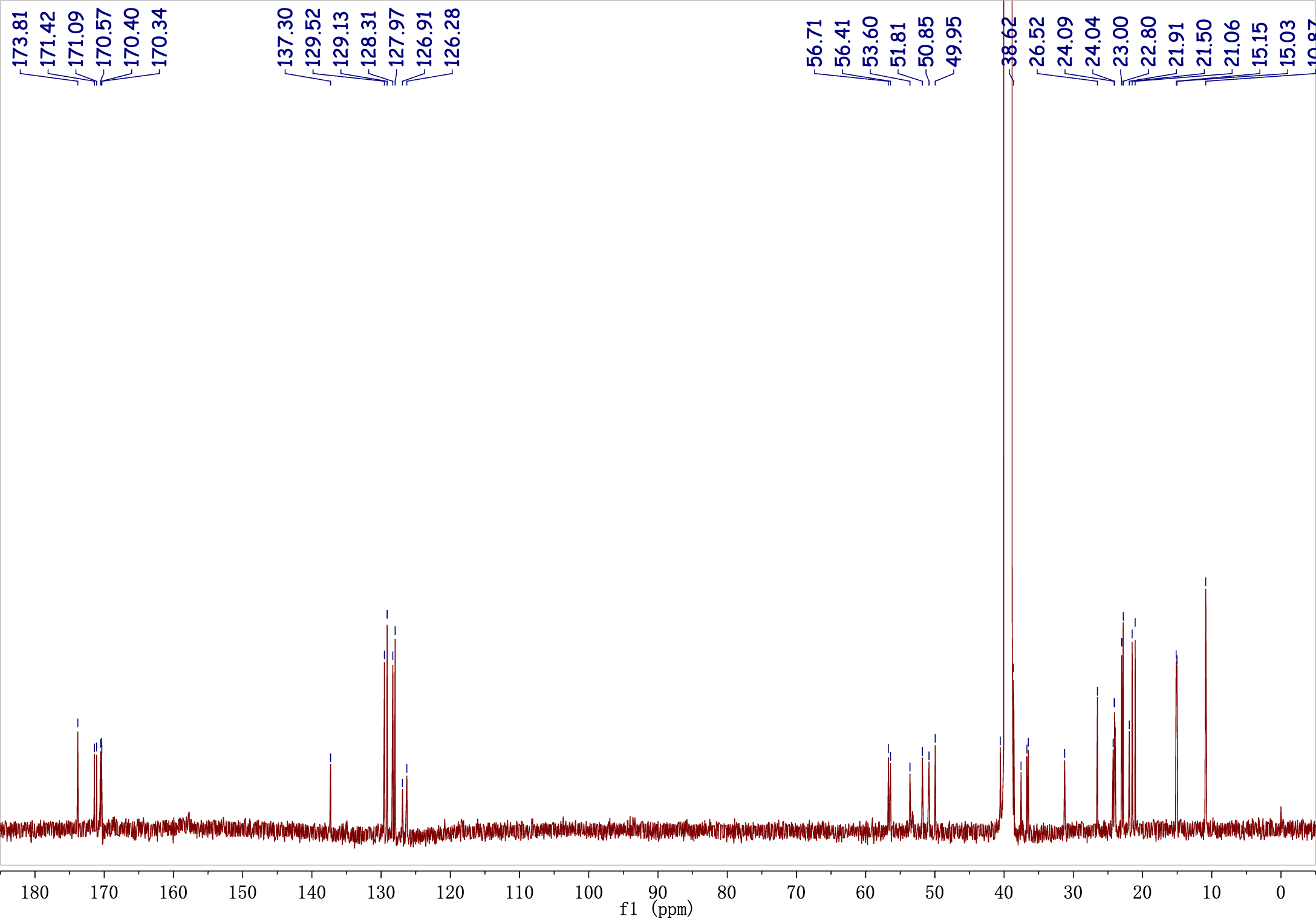
